# Enhancing top-down proteomics of brain tissue with FAIMS

**DOI:** 10.1101/2021.01.18.427216

**Authors:** James M. Fulcher, Aman Makaju, Ronald J. Moore, Mowei Zhou, David A. Bennett, Philip L. De Jager, Wei-Jun Qian, Ljiljana Paša-Tolić, Vladislav A. Petyuk

**Affiliations:** Biological Sciences Division, Pacific Northwest National Laboratory, Richland, WA 99352, USA; Life Sciences Mass Spectrometry Unit, Thermo Fisher Scientific, San Jose, CA 95134, USA; Environmental Molecular Sciences Laboratory, Pacific Northwest National Laboratory, Richland, Washington 99354, USA; Rush Alzheimer’s Disease Center, Rush University Medical Center, Chicago, IL 60612, USA; Department of Neurology, Center for Translational & Computational Neuroimmunology, Columbia University Medical Center, New York, NY 10032, USA

**Keywords:** FAIMS, differential mobility spectrometry, top-down proteomics, Alzheimer’s, brain tissue, ion mobility

## Abstract

Proteomic investigations of Alzheimer’s and Parkinson’s disease have provided valuable insights into neurodegenerative disorders. Thus far, these investigations have largely been restricted to bottom-up approaches, hindering the degree to which one can characterize a protein’s “intact” state. Top-down proteomics (TDP) overcomes this limitation, however it is typically limited to observing only the most abundant proteoforms and of a relatively small size. Therefore, offline fractionation techniques are commonly used to reduce sample complexity, limiting throughput. A higher throughput alternative is online fractionation, such as gas phase high-field asymmetric waveform ion mobility spectrometry (FAIMS). Utilizing a high complexity sample derived from Alzheimer’s disease brain tissue, we describe how the addition of FAIMS to TDP can robustly improve the depth of proteome coverage. For example, implementation of FAIMS at −50 compensation voltage (CV) more than doubled the mean number of non-redundant proteoforms observed (1,833 ± 17, n = 3), compared to without (754 ± 35 proteoforms). We also found FAIMS can influence the transmission of proteoforms and their charge envelopes based on their size. Importantly, FAIMS enabled the identification of intact amyloid beta (Aβ) proteoforms, including the aggregation-prone Aβ_1-42_ variant which is strongly linked to Alzheimer’s disease.

## Introduction

Over the last two decades, the incidence of neurodegenerative diseases has more than doubled worldwide, with Alzheimer’s disease (AD) as the most prevalent form.^1^ Two protein species, amyloid beta (Aβ) peptides and phosphorylated microtubule-associated protein tau (tau),^2–5^ have been found to be strongly associated with AD. Therefore, detailed proteome characterization of AD has been of importance. ^6, 7^ Mass spectrometry has played a central role in these investigations,^8^ particularly bottom-up approaches.^9–16^ However, the protease digestion required for bottom-up analyses impedes the capturing of a protein’s complete state, which can vary due to genetic alleles, alternative splicing, proteolytic processing, and post-translational modifications (referred to as “proteoforms”). ^17–20^ Since top-down proteomic (TDP) approaches analyze proteins in an intact state, the likelihood of capturing proteoforms associated with certain pathologies is greater and allows for a stronger, more direct connection between genotype and phenotype.^20–22^ For example, TDP is particularly well suited for capturing endogenous proteolytic fragments derived from proteins such as tau, which have been linked to Alzheimer’s pathology.^23–26^ However, TDP of complex samples requires offline fractionation techniques to reduce sample complexity, as the human proteome spans several orders of magnitude in size and abundance, and proteins are generally not well-resolved with reverse-phase high-performance liquid chromatography (RP-HPLC).^27^ This offline fractionation can be accomplished with gel-eluted liquid fraction entrapment electrophoresis (GELFrEE), size-exclusion chromatography (SEC), ion exchange chromatography, or affinity purification, to name a few.^22, 28–31^ Unfortunately, offline fractionation typically reduces throughput.^32^ An attractive online fractionation alternative that can be introduced between the LC and MS dimensions with no impact on throughput is gas-phase ion mobility separation. High field asymmetric waveform ion mobility spectrometry (FAIMS) is particularly well suited to this task with the recent introduction of the FAIMS Pro device, whose modular design allows facile incorporation of ion mobility onto several current Orbitrap instruments. ^33^ FAIMS, also referred to as differential mobility spectrometry (DMS), separates ions in a carrier gas based on combinations of factors such as size, charge, or shape through the introduction of an asymmetric waveform with high and low electric fields.^34^ To prevent collision of the ions with the electrode, a deviation in the ion’s path is introduced through the application of a DC compensation voltage (CV), ^35^ allowing selective transmission of that ion.^34^ The application of FAIMS to intact protein analysis has largely been restricted to the separation of conformers of individual proteins or small combinations of proteins,^36–41^ however, FAIMS has recently been applied to intact and native protein analyses using liquid extraction surface analysis (LESA).^42–46^ Here we apply FAIMS-TDP analysis to a whole-tissue sample from the medial frontal cortex (MFC) of an Alzheimer’s patient. Scanning across a CV range of −50 to −20 CV with external stepping allowed us to determine how modulation of FAIMS CV influences the characteristics of the ions transmitted through the cylindrical FAIMS unit, and how this relationship can be exploited to target proteoforms based on size. FAIMS-TDP more than doubled proteoform identifications at a single CV compared to without FAIMS and enabled deeper interrogation of proteoforms relevant to neurodegenerative diseases, including α-, β-, and γ-synucleins, PARK7, tau splice isoforms, and several intact Aβ proteoforms. Taken together, this work describes how greater identifications of proteoforms and proteome coverage can be achieved reproducibly via gas-phase fractionation with FAIMS in TDP.

## Methods

### Sample Preparation

Medial frontal cortex (MFC) sample was received from Rush University

Alzheimer’s Disease Center. The patient’s overall cognitive diagnostic category was Alzheimer’s dementia with no other causes of cognitive impairment,^47^ while the postmortem interval was 230 minutes. 33 mg of MFC tissue stored at −80°C was transferred to 1.5 mL LoBind Eppendorf tubes (Eppendorf, Cat# 022431081) and combined with 0.5 mL homogenization buffer consisting of 8 M urea, 10 mM ammonium bicarbonate (ABC) and 10 mM tris(2-carboxyethyl)phosphine (TCEP). Homogenization was performed with a handheld pellet pestle (BioVortexer with SpiralPestle; Biospec 1083 and 1017) before combination with another 0.5 mL homogenization buffer. The sample was then incubated at 37°C for 60 minutes with 1,200 RPM mixing using a ThermoMixer to allow for extraction and denaturation of cellular proteins. Pelleting of urea-insoluble material was accomplished with centrifugation at 18,000 *g* for 20 minutes at 22°C and the resulting supernatant was then added to 3 mL wash buffer (WB; 8 M Urea, 10 mm ABC) within a 4 mL 100 kDa MWCO filter to remove large MW species. After 1 hr of centrifugation at 22°C and 5,000 *g*, the retentate volume was 200 µL and filtrate 3.8 mL. The 3.8 mL filtrate was transferred to a 3 kDa MWCO filter to remove small molecule contaminants and low MW peptides. The 3 kDa MWCO filter was centrifuged at 22°C and 7,300 *g* for 1 hr, giving a retentate volume of 200 µL. The 200 µL retentate from the 100 kDa MWCO filter was washed once more by diluting to 4 mL with WB and filtering it again through the same filter. This flowthrough was added to the same 3 kDa MWCO filter as above and concentrated one more time. The final 200 µL retentate from the 3 kDa MWCO filter wash was transferred to a 1.5 mL LoBind Eppendorf tube. To acidify the sample, 10 µL of 10% formic acid (FA) was added, resulting in 0.5% FA concentration in the final sample. The sample was centrifuged at 18,000*g* for 15 minutes to remove precipitated material before measuring the protein concentration using BCA assay. To account for any potential buffer contributions to the BCA assay, BSA standards were prepared in 8 M urea, 10 mM ABC, 0.5% FA. Protein concentration by BCA assay was determined to be 1.1 mg/mL, corresponding to a final yield of 0.22 mg total recovered protein in the ~200 µL sample. Sample was diluted to 0.5 mg/mL with WB containing 0.5% FA and stored at −80°C until LC-MS analysis.

### LC-MS/MS Analysis and FAIMS Settings

Samples were analyzed using a Waters NanoACQUITY UPLC system with mobile phases consisting of 0.2% FA in H_2_O (Mobile Phase A) and 0.2% FA in ACN (Mobile Phase B). Both trapping-precolumn (100 µm i.d., 5-cm length) and analytical column (75 µm i.d., 50-cm length) were slurry-packed with C2 packing material (5 µm and 3 µm for trap/analytical respectively, 300 Å, Separation Methods Technology). Samples were loaded into a 5 µL loop, corresponding to 2.5 µg of loaded material amount, and injected onto the trapping column with an isocratic flow of 5% B at 3 µL/min over 20 min. Separation was performed with a 5% to 50% B gradient over 180 min at 300 nL/min. For MS/MS analysis of proteins, the NanoACQUITY system was coupled to a Thermo Scientific™ Orbitrap Fusion™ Lumos™ Tribrid™ mass spectrometer equipped with the FAIMS Pro interface. Source parameters included electrospray voltage of 2.2 kV, transfer capillary temperature of 275°C, and ion funnel RF amplitude of 60%. FAIMS was set to standard resolution without supplementary user-controlled carrier gas flow and a dispersion voltage (DV) of −5 kV (equivalent to a dispersion field of −33.3 kV/cm),^46^ while the compensation voltage (CV) was varied depending on the experiment. The Fusion Lumos was set to “intact protein” application mode, which lowers the HCD cell N2 pressure to 2 mTorr, and data was collected as full profile. MS^1^ and MS^2^ data were acquired at a resolution of 120k and 60k, microscans of 3 and 2, across a 500-2,000 *m/z* and 400-2,000 *m/z* range, and with AGC targets of 5E6 and 5E5, respectively. MS^1^ and MS^2^ were acquired with a maximum inject time of 400 ms as well. Data dependent settings included selection of top 6 most intense ions, exclusion of ions lower than charge state 5+, inclusion of undetermined charge states, and dynamic exclusion after 1 observation for 30 s. Ions selected for MS^2^ were isolated over a +/− 1.5 *m/z* window and fragmented through collision-induced dissociation (CID) with a collision energy of 35%. Datasets for each FAIMS CV were collected in triplicate. All raw files have been deposited into the MassIVE data repository, and can be accessed via accession MSV000086696, or alternatively with PXD023607 on ProteomeXchange.

## Data Analysis

Proteoform identification was performed with TopPIC version 1.3.^48^ Settings for TopPIC included a precursor window of 3 *m/z* (to account for isotopic envelope), mass error tolerance of 15 ppm, a max/min unknown mass shift of 500 Da and a maximum number of allowed unknown modifications of 1. MS^2^ spectra were searched against a database concatenated with entries from *Homo sapiens* Swiss-Prot (20,352), Swiss-Prot splice variants (22,000), and TrEMBL (54,436), as well as common contaminants. Identified proteoforms were filtered to an FDR of 1% through TopPIC. Downstream data analysis was performed in the R environment for statistical computing and figure generation.^49^ In the downstream analysis, proteoforms from the same gene with the same starting and ending amino acids that were found to be within +/− 5 Da were combined as a single proteoform in order to increase the stringency of our assignments. Although the 5 Da threshold is arbitrary, this approach compensated for incorrectly assigned monoisotopic peaks, ambiguity in unknown mass shifts, and other artifactual deviations. In order to determine the relative standard deviation of proteoforms within replicates, we utilized the ‘feature intensity’ output from TopPIC. Fragmentation sequence coverage maps and associated spectra were generated using LCMsSpectator.^50^ Graphical abstract and image assets within Figure 1 were created using Biorender.com.

**Figure 1:**
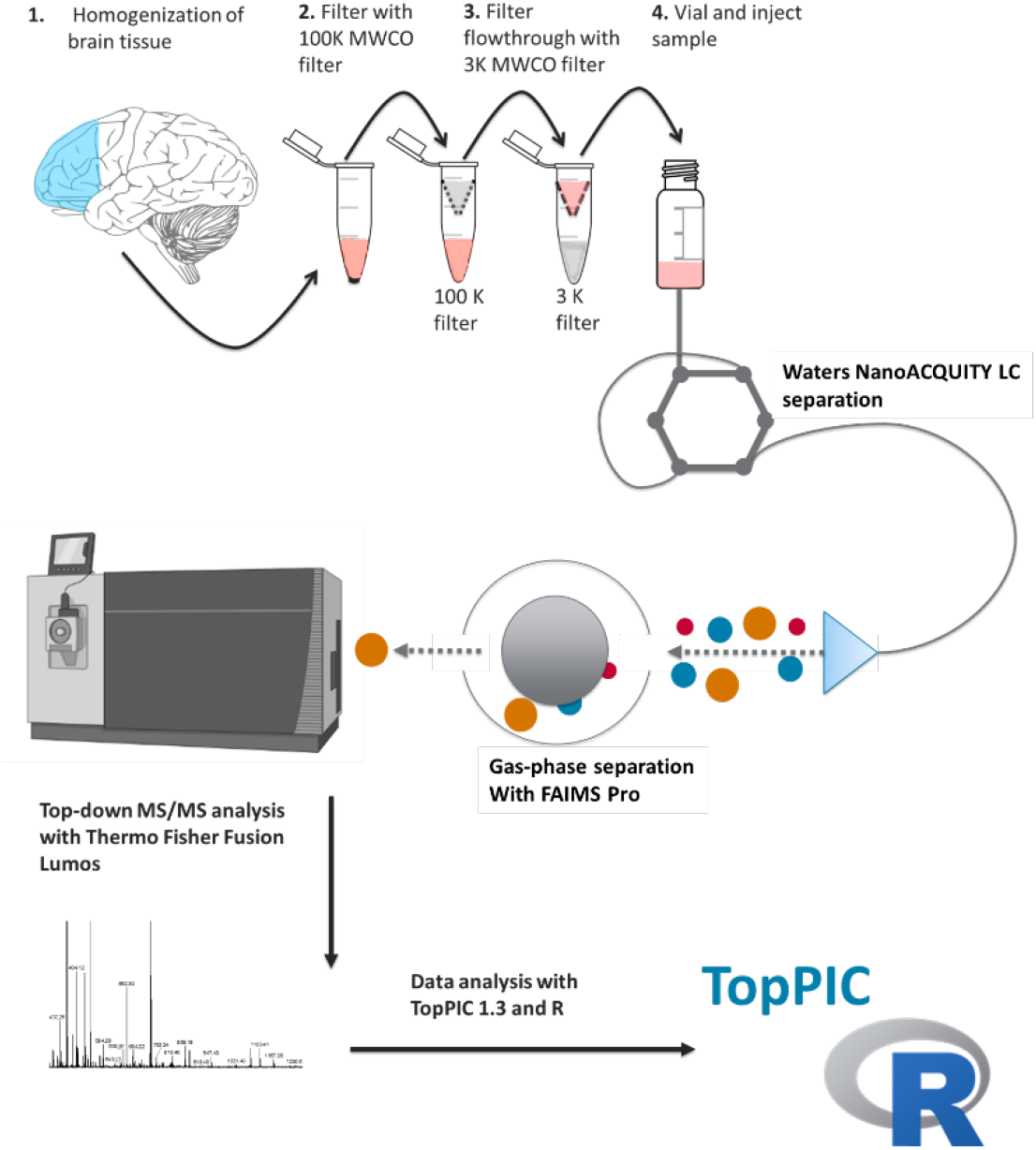
Workflow demonstrating sample preparation, LC-MS, FAIMS-TDP, and data analysis with TopPic/R.

## Results and Discussion

### Addition of FAIMS Robustly Increases Proteome Coverage

The cortex tissue sample from a patient diagnosed with Alzheimer’s was processed following the workflow in **Figure 1**.

We first sought to analyze the performance of FAIMS at various CVs in comparison to without FAIMS (referred to as “No FAIMS”). It is worth remarking that the FAIMS interface produces a modest loss of ion transmission, presumably due to the longer ion path through the source.^51^ Therefore, to better represent typical conditions and ensure equitable comparisons, the “No FAIMS” data were collected using the same sample without the FAIMS unit installed. These “No FAIMS” data were collected in triplicate and analyzed according to the workflow in **Figure 1** with a top 6 DDA MS^2^ analysis.

The “No FAIMS” datasets on average identified 754 ± 35 proteoforms (**Figure 2A** and **Table S1**) and collectively identified 1,073 unique (non-redundant) proteoforms (**Figure 2B**) derived from 293 unique genes (**Figure 2C**), covering 29,359 amino acids across the proteome (**Figure 2D**). The metric “proteome coverage” (**Figure 2D**) is defined as non-redundant amino acids covered by each proteoform’s sequence. Essentially, this metric accounts for both the length and diversity of the identified proteoforms. We consider this to be a more balanced metric compared to the raw number of proteoforms and genes, as it avoids biases towards smaller proteoforms. Although simple counts of proteoforms or genes are intuitive, these metrics appear to be strongly influenced by short proteolytic fragments.

**Figure 2:**
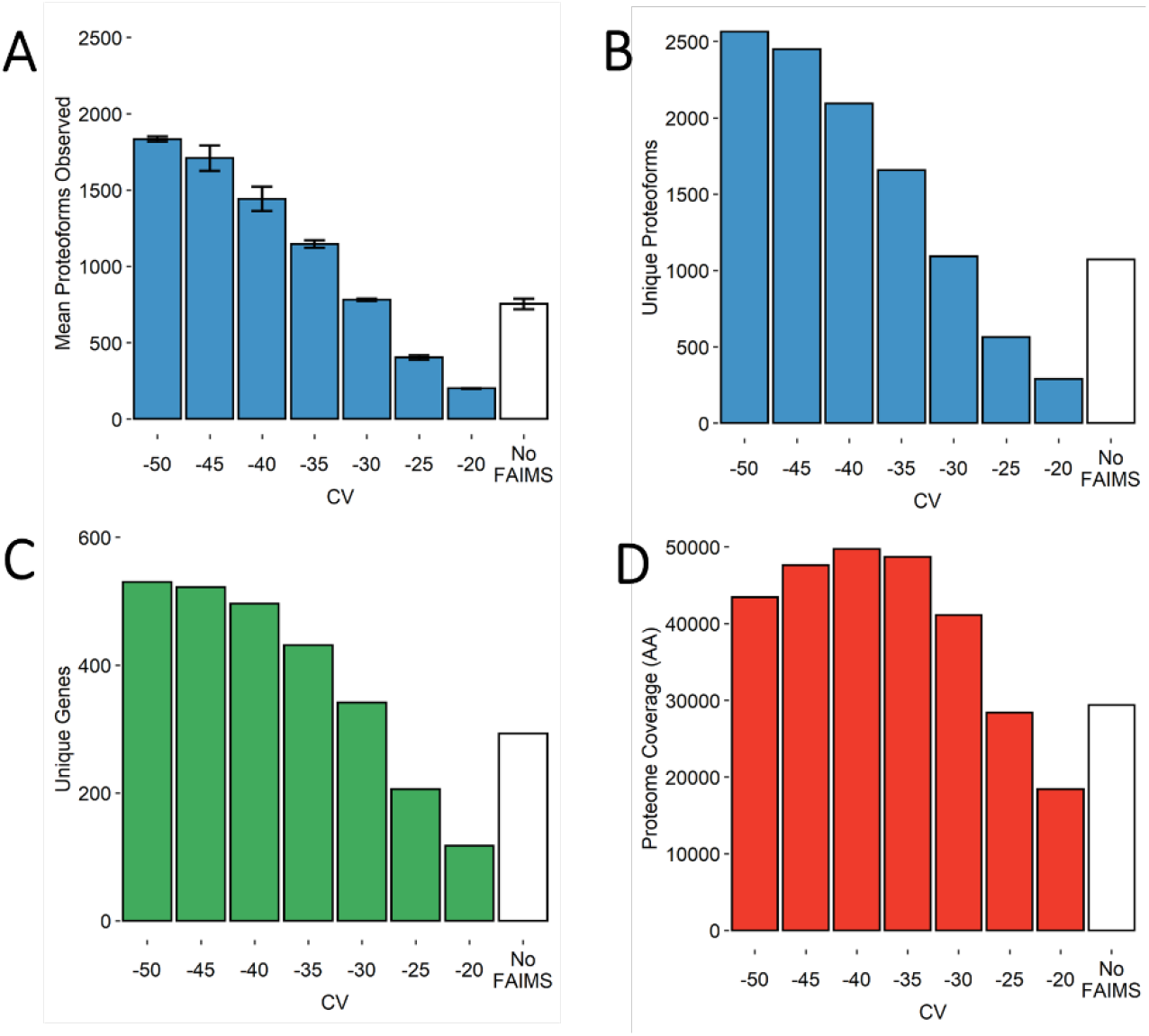
Bar charts demonstrating several metrics used in the comparison of FAIMS CVs in steps of 5V with “No FAIMS” datasets. (**A**) Mean number of proteoform identifications per replicate (n = 3). Error bars represent standard deviation from the mean. (**B**) Total unique proteoforms found across all three replicates for each FAIMS CV or “No FAIMS”. (**C**) Total unique genes found across all three replicates for each FAIMS CV or “No FAIMS. (**D**) Total proteome coverage, or non-redundant amino acids covered by every proteoform’s sequence, found across all three replicates for each FAIMS CV or “No FAIMS.

The FAIMS datasets were collected from −50 to −20 CV scanned in steps of 5 V. Each of the resulting 7 CVs were collected in triplicate. With respect to proteoforms and genes, FAIMS outperformed “No FAIMS” for all the CVs tested within the −50 to −30 CV range (**Figure 2A-C**). For example, the 3 datasets at −50 CV identified 1,833 ± 17 proteoforms, an average increase of ~140% per run compared to “No FAIMS”. Collectively, these three datasets at −50 CV identified 2,564 unique proteoforms from 530 unique genes covering a total of 43,437 amino acids, an increase of 95%, 69%, and 69% over the three “No FAIMS” datasets, respectively (**Figure 2B-D** and **Table S1**). Also, although FAIMS at −25 V observes fewer unique proteoforms and genes compared to “No FAIMS”, the proteoforms themselves are much longer on average. Thus, when considering proteoform length, (**Figure 2D**), the −25 V datasets covers nearly as many amino acids from the proteome (28,398) relative to “No FAIMS” (29,359), doing so with only half as many proteoforms (**Figure 2B**).

We also investigated if FAIMS could provide as robust and reproducible an analysis compared to without FAIMS. We assessed quantification reproducibility by calculating the relative standard deviation (RSD, equivalent to coefficient of variation) of each proteoform’s feature intensity, provided the proteoform met the criterion of being observed across all three replicate datasets for each CV setting. Because the FAIMS and “No FAIMS” datasets were all collected from a single biological sample, variance among the replicates should be exclusively attributed to instrumental factors. The boxplot in **Figure 3** (as well as individual histograms in **Figure S1**) demonstrates how the distributions of RSDs compare between the FAIMS and “No FAIMS” datasets.

**Figure 3:**
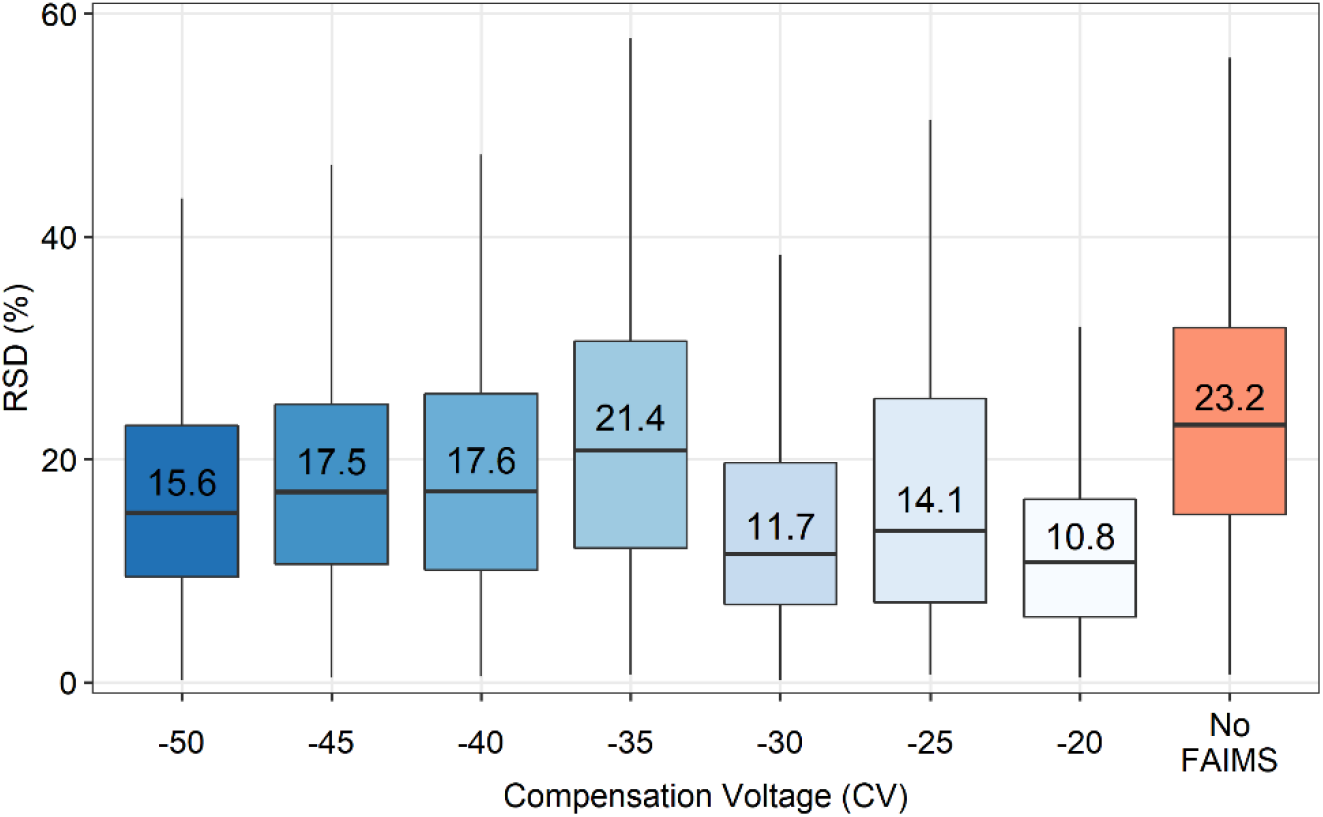
Boxplot displaying the distributions of the relative standard deviations (RSDs) determined from the feature intensities of proteoforms found in FAIMS and “No FAIMS” replicates. The median RSD for each condition is written within the upper box, above the line indicating its position. The upper and lower hinges correspond to the first and third quartiles (25^th^ and 75^th^ percentiles), while the upper and lower whiskers represent 1.5 times the interquartile range (IQR). Outliers beyond the whiskers are not plotted.

Based on median RSD, addition of FAIMS resulted in similar or slightly improved quantification quality relative to “No FAIMS”. The improvement in RSDs among lower voltages (−20 to −30 CV) may be related to the observation that fewer proteoforms are transmitted within that range, providing higher quality MS^1^ spectra and therefore better estimations of feature intensity. Furthermore, it is worth noting that the RSDs determined from these replicates are similar to previous top-down^50^ and bottom-up^52, 53^ experiments performed with similar instrumentation. Taken together, this data demonstrates that the addition of FAIMS to TDP robustly increases proteoform identifications, without any sacrifice of quantification quality.

We next investigated the overlap in identifications between the CVs, and which combination of CVs could provide the largest non-redundant set of observations. Using overlap coefficients, defined as the intersection between two sets divided by the smaller of the two sets, we determined the similarity between each dataset. The overall mean overlap coefficient for both proteoform and gene identifications within each FAIMS CV was 0.76. This is comparable to overlap coefficient from “No FAIMS” datasets (0.77), suggesting that the technical replicates show similar reproducibility. A heatmap displaying the FAIMS overlap coefficients of proteoforms and genes is shown in **Figure 4** and **Figure S2**, respectively.

**Figure 4:**
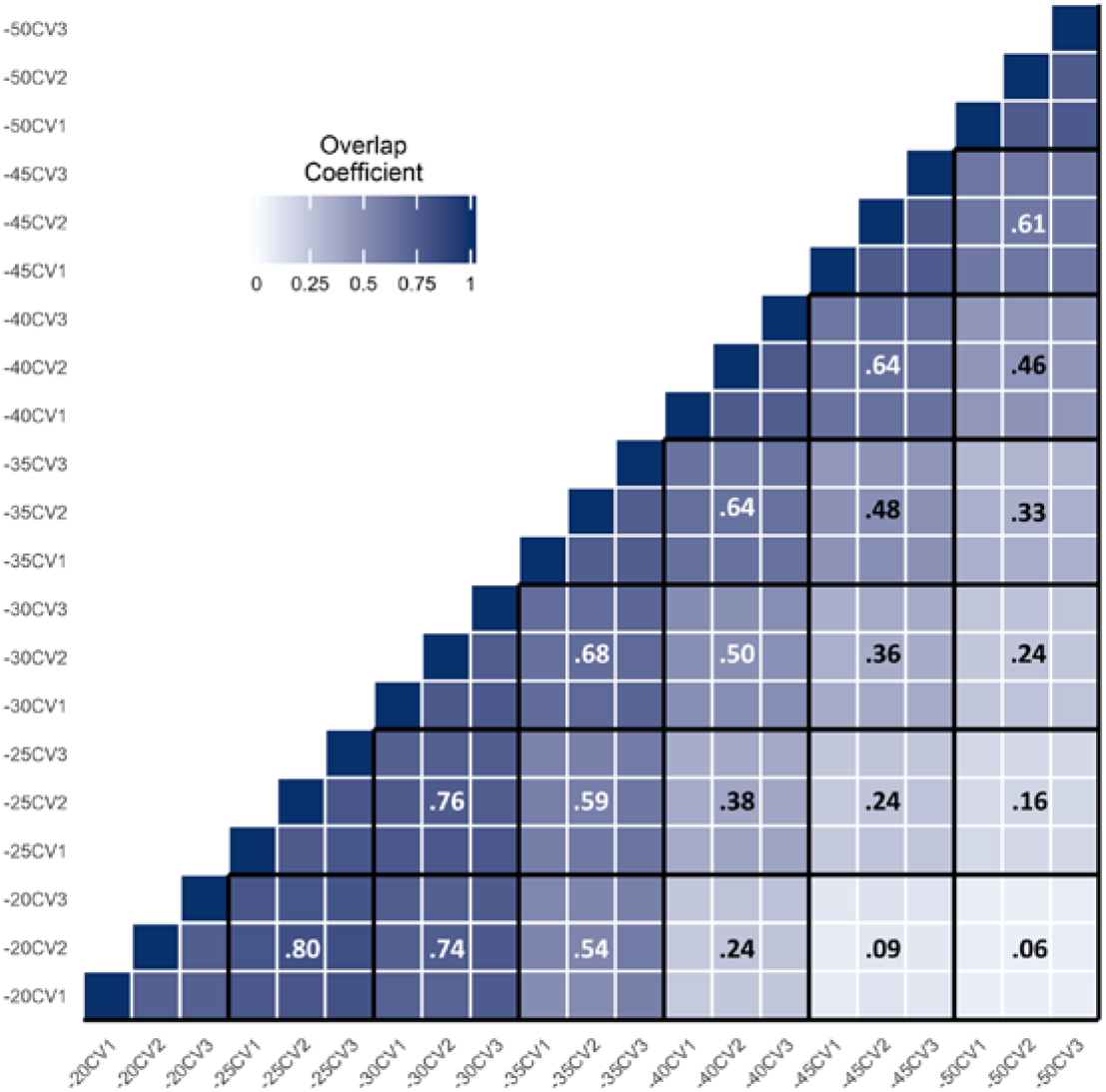
Heatmap generated in R comparing the overlap coefficients of each CV replicate based on proteoforms identified with TopPIC. White lines separate replicates while black lines separate different CVs. Mean overlap coefficients of each CV’s replicates are shown in the middle block of the CVs being compared. White text color is used on overlap coefficients >= 0.5 and black text color < 0.5 to improve visibility.

Notably, the overlap of gene identifications across the CV space is much greater relative to proteoforms, although it is particularly noticeable when comparing the longest CV distances. For example, the overlap between −50 and −20 CV is 0.64 at the gene level and 0.06 at the proteoform level. This was expected, as each gene can potentially be represented by multiple proteoforms. **Figure 4** demonstrates that while a 5 to 10 V difference in CV produces overlap coefficients mostly above 0.5, distances greater than 15 V produce larger degrees of dissimilarity and are therefore better spaced for capturing sufficiently different sets of proteoforms. With these datasets, we next pursued determining which combinations of CVs are optimal for achieving maximal gene identifications, proteoform identifications, and sequence coverage with constraints on the number of CVs for each combination. First, we established the optimal combinations for any number of CVs from 1 to 7 using the metrics unique proteoforms, genes, and proteome sequence coverage (**Figure 5**).

**Figure 5:**
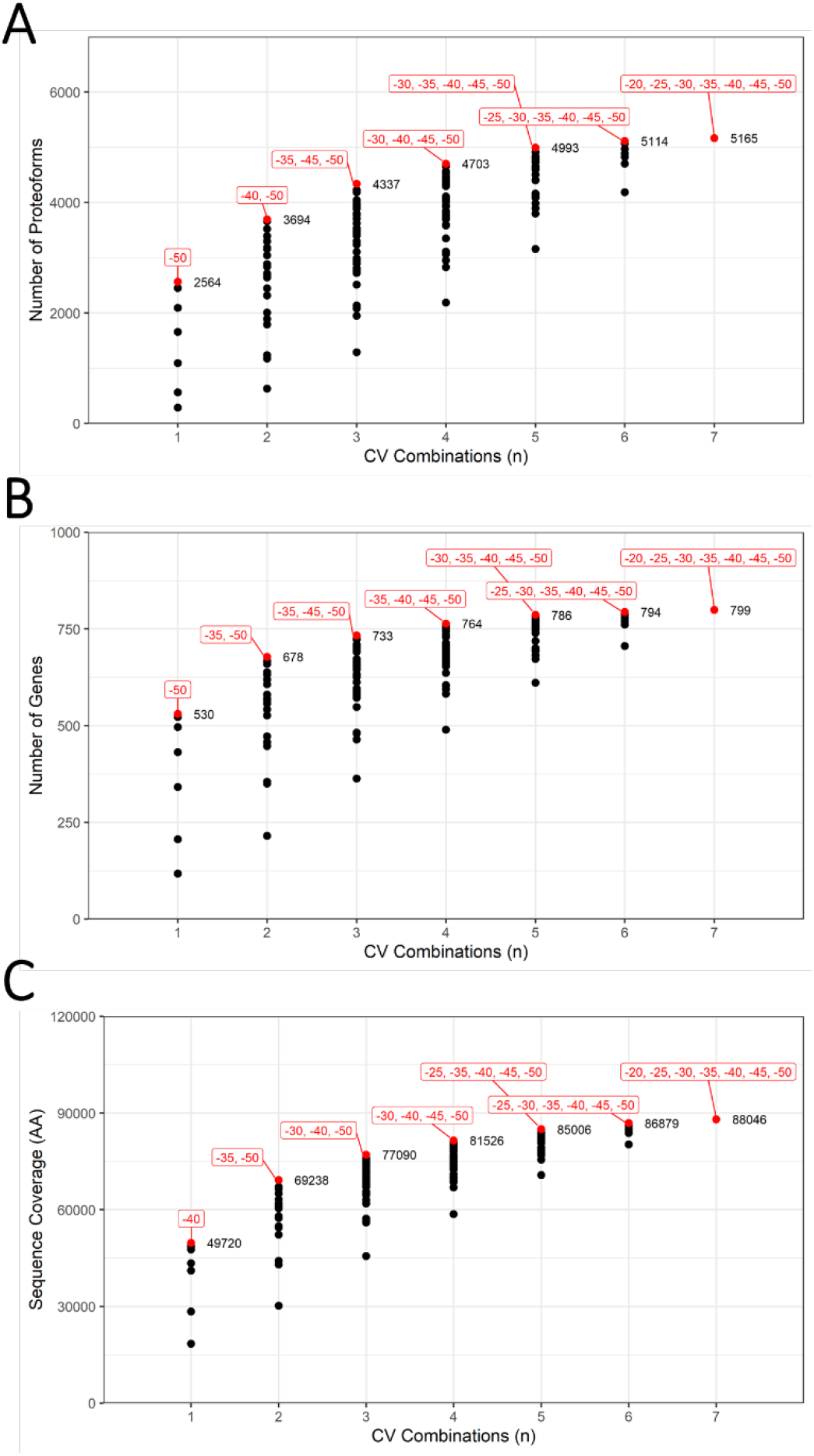
Combinatorial analysis of FAIMS CVs with regards to (**A**) number of proteoforms, (**B**) number of genes, and (**C**) sequence coverage. For each combination, the highest number plotted is highlighted in red and labeled with the CVs for that specific combination.

Using combinations limited to 3 CVs as an example, **Figure 5A and 5B** detail how −35, −40, and −50 V are ideal if one desires to maximize the number of proteoforms or genes. The proteoforms and genes identified in these 3 CVs (9 datasets) cover 84% of the total unique proteoforms and 92% of the total genes over the entire set of 21 datasets. However, when weighing a proteoform’s length with sequence coverage as a metric, the ideal combination is at −30, −40 and −50 CV, covering 88% of the unique amino acids from all 21 FAIMS datasets with only a modest loss of proteoform and gene identifications (4221 proteoforms and 723 genes, **Figure 5C**). Considering these results with the overlap analysis (**Figure 4**), 15 V or greater distances between CVs provide the least amount of overlap between proteoforms, while 5-10 V separation within the −50 to −30 CV maximizes genes, proteoforms, and proteome sequence coverage.

To further determine the impact of FAIMS on the depth of proteome coverage, we compared the genes identified from our top-down data sets to a typical bottom-up analysis of human brain. Comprehensive bottom-up datasets of human brain tissue allowed us to estimate abundances for 8,528 proteins using weighted spectral counting.^54–56^ Proteins were binned into 10 different abundance percentiles based on spectral counts compiled from several bottom-up datasets of the human brain. By cross-referencing with our top-down datasets, we were able to determine where the genes, and the proteoforms they are derived from, rank in terms of estimated protein abundance (using dataset-normalized spectral counts) in the human brain (**Figure 6**).

**Figure 6:**
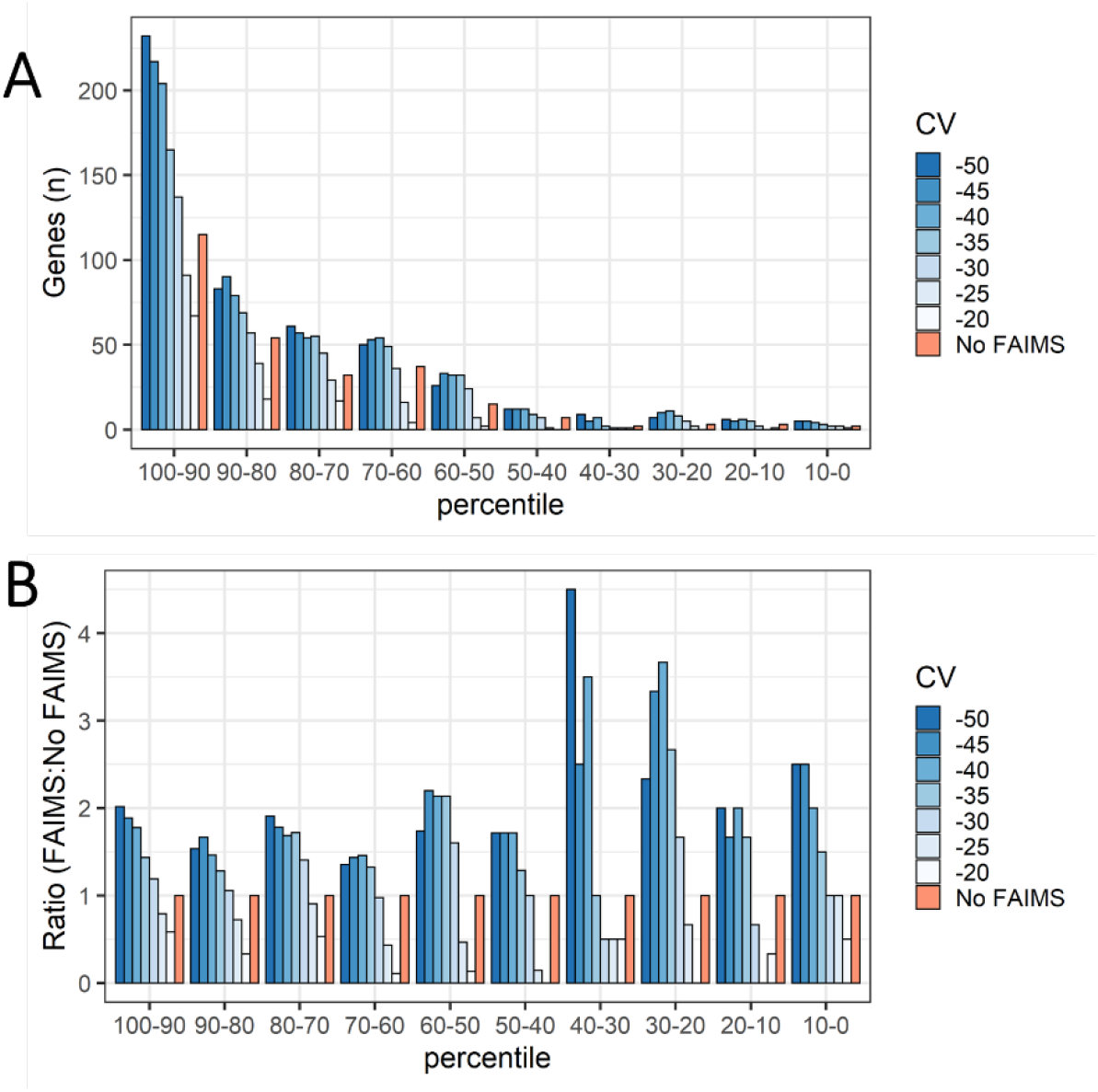
Bottom-up datasests from human brain tissue were used to estimate protein abundance by weighted spectral counting. These proteins were then binned into 10 different abundance percentiles. (**A**) Comparison between the genes found in FAIMS and “No FAIMS” datasets across the 10 different abundance percentiles. (**B**) The number of genes for each FAIMS CV were normalized to the number of genes found in the “No FAIMS” datasets for each abundance percentile.

One immediate and apparent observation is that the higher the abundance percentile, the higher the probability of identification with TDP. About half of the genes identified from the top-down datasets (with or without FAIMS) were contained within the top 80^th^ percentile of abundance (**Figure 6A**). As anticipated, we found that FAIMS could improve identifications in lower-abundant percentiles over “No FAIMS”, increasing the depth of the observable proteome. By determining the ratio of FAIMS to “No FAIMS” across all percentiles, an appreciable increase in identifications below the 40^th^ percentile is apparent within CVs in the −40 to −50 range (**Figure 6B**). However, when considering the absolute number of genes, the major contribution of FAIMS’ advantage is through broadening the spectrum of identifications across all abundance percentiles.

### Relationship between FAIMS Compensation Voltage and Transmission of Proteoforms

With FAIMS, we also noted a trend between FAIMS CV and proteoform molecular weight (**Figure 7**).

**Figure 7:**
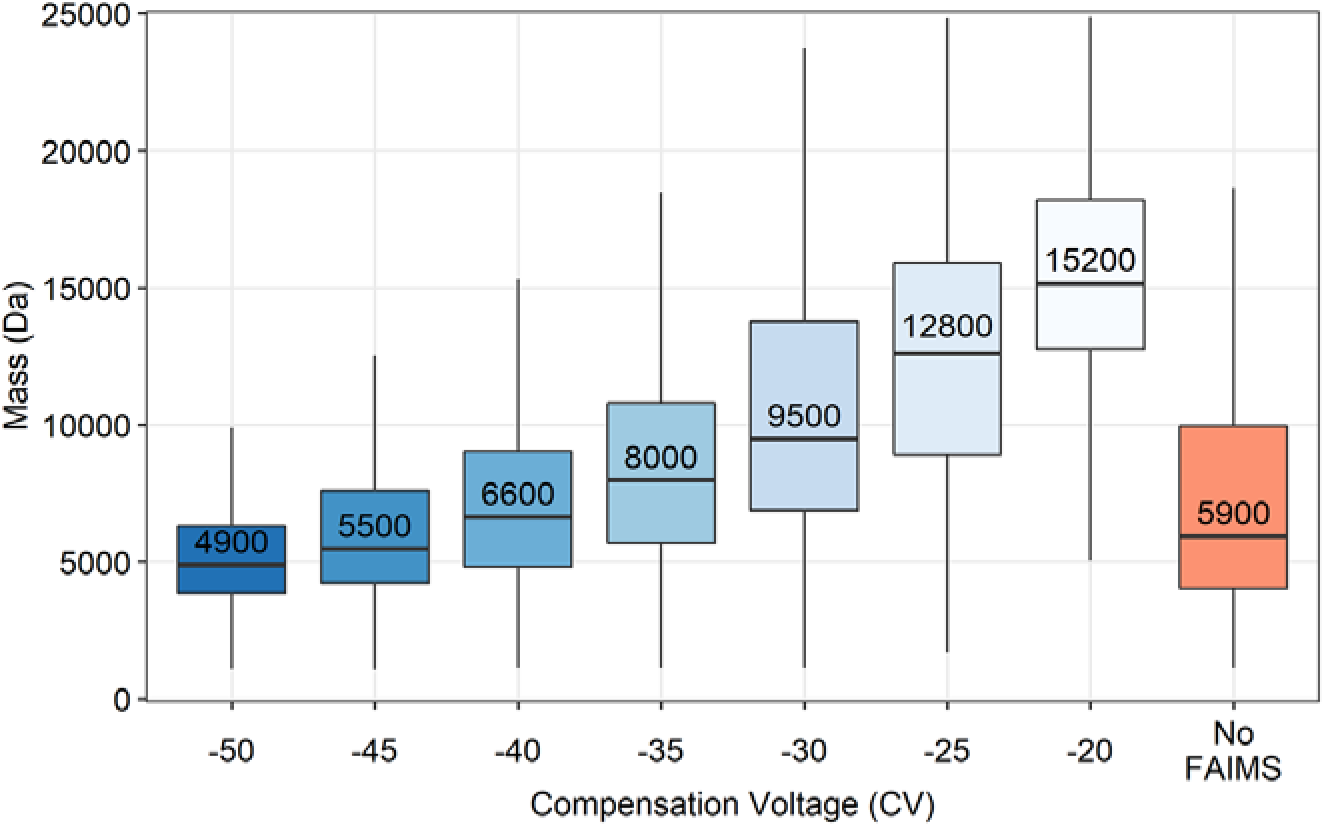
Boxplot displaying distributions of proteoform masses identified with TopPic against tested FAIMS CVs. Each distribution includes only non-redundant proteoforms found within all three replicates. The median proteoform mass for each condition is written within the upper box, above the line indicating its position. The upper and lower hinges correspond to the first and third quartiles (25^th^ and 75^th^ percentiles), while the upper and lower whiskers represent 1.5 times the interquartile range (IQR). Outliers beyond the whiskers are not plotted.

At −50 CV the median proteoform mass is ~5 kDa and increases to ~15 kDa at −20 CV. Based on these mass distributions (**Figure 7**), CVs less than −50 V appear well suited for top-down or middle-down proteomic experiments while CVs greater than −50 V may best benefit peptidomic or bottom-up experiments. Interestingly, although the majority of spectra identifying a single proteoform were limited to being found within a 10 V range (3,599 out of 5,165), there were 7 proteoforms observed across the entire CV range from −50 to −20. Presumably, this can be attributed to their high abundance in the sample, particularly in the case of ubiquitin (UBB), myelin basic protein (MBP), and acyl-CoA-binding protein (ACBP). However, it is also likely that the different charge states of these proteoforms adopt several gas-phase conformations depending on their proton-isomerization, impacting their mobilities.^38^ These 7 proteoforms allowed us to investigate how the charge state envelope is differentially transmitted through the modulation of CV. As a proxy for the charge state distribution at each CV, we used the median charge state based on the PrSMs identified at each CV and tracked how this value changed as a function of CV starting at −50 CV. The MS^1^ scans in **Figure S3** demonstrate how the median PrSM charge state tracks the charge state distribution of UBB. Of these 7 proteoforms, 4 followed an inverse relationship, with median observed charge state increasing as CV was decreased (**Figure 8**).

**Figure 8:**
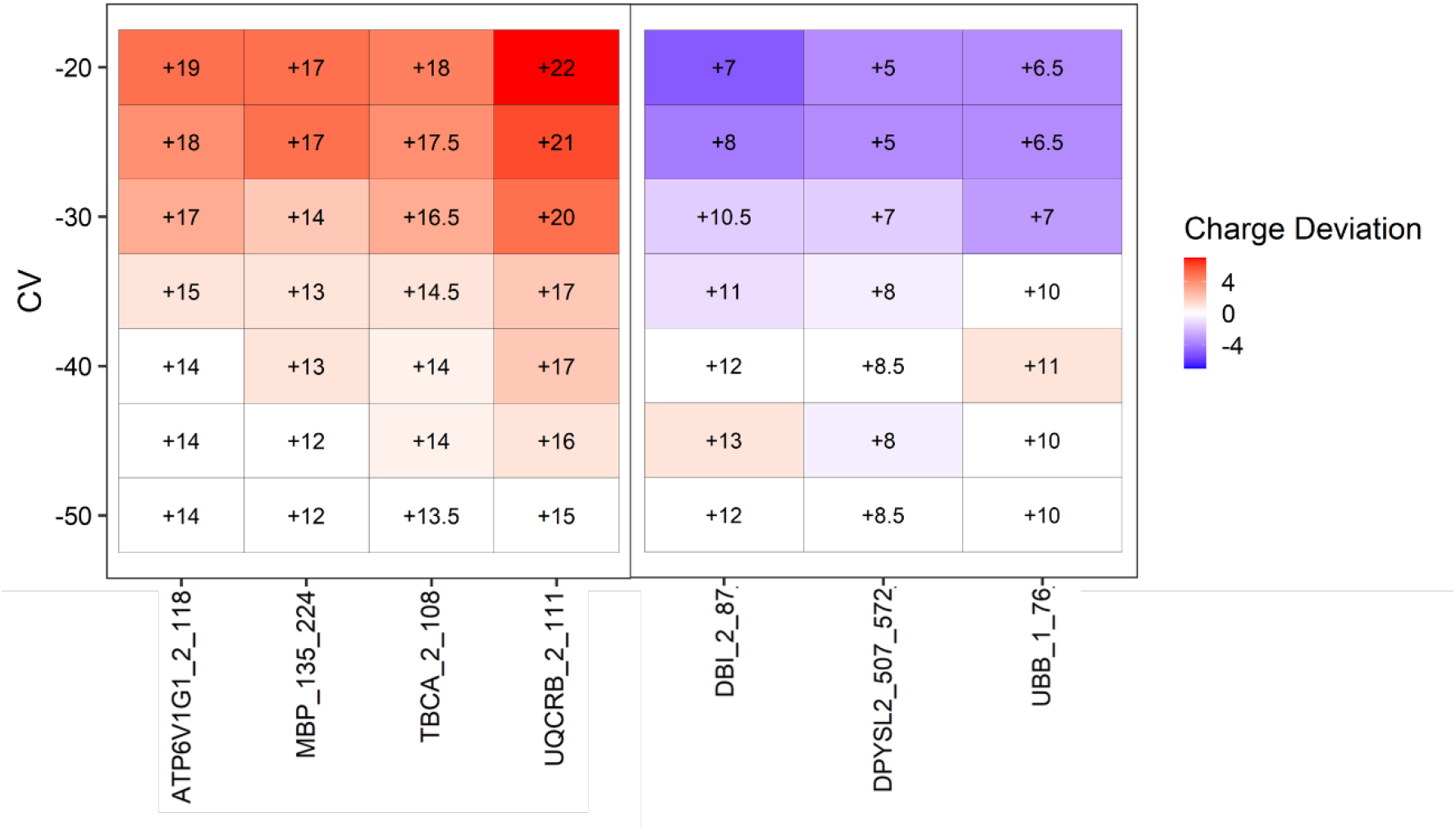
Heatmap of the median PrSM charge states from 7 proteoforms identified across the −50 to −20 CV range, filtered by >= 4 PrSMs in order to increase the confidence in identifications and provide a reasonable estimate of median charge state. The median PrSM charge state is shown within each block, while the deviation of the PrSM charge state is indicated in red (increasing) or blue (decreasing) relative to −50 CV. Proteoforms on the x-axis are written as the gene name, starting amino acid, and ending amino acid.

It’s worth noting this trend has generally been observed with peptides and small proteins.^51, 57–59^ Surprisingly, 3 proteoforms showed the opposite relationship where decreasing CV decreased the median charge states observed (**Figure 8**). A cursory examination of the mean precursor mass between the two groups suggests that larger precursors are more likely to favor higher charge states as CV is decreased. To validate these relationships further, we expanded this analysis to include proteoforms that were identified within a more modest, but still wide, 20-30 CV range (n = 256 proteoforms). Here, proteoforms whose median charge shifted greater than 1 charge across the entire CV range were binned into the two different groups depending on the direction of that shift. Those that shifted less than one charge were considered “neutral” with respect to changing CV. Similar to the previous results, larger precursors were significantly more likely to be associated with an inverse relationship between observed charge states and CV (**Table S2**). With decreasing CV, the majority of proteoforms appeared to transmit at lower charge states (n = 177) compared to those that favored higher charge states (n = 39). The remainder (n=40) were considered “neutral” with respect to changes in CV. Proteoforms binned within the neutral group demonstrated an average precursor mass between the inverse and direct groups, once again suggesting the mass of the proteoform is intrinsically linked to this behavior. Other primary sequence-based parameters such as basic/acidic amino acid composition and aliphatic index were not significantly correlated however (**Table S2**). Although the proteins being introduced into the gas-phase are presumably denatured, factors typically associated with native proteins such as dipole moment or collisional-cross section may have better predictive value towards this phenomenon.^59–61^ Our results demonstrate how CV can be used to filter proteins of different masses, and how a protein’s charge state envelope can be differentially transmitted through FAIMS as well.

### Utility of Protein Fragments in Top-Down Proteomic Experiments

Surprisingly, a significant number of proteoforms we identified were fragments of larger proteins. We found that only 25% of the unique proteoforms identified covered greater than half of the protein’s sequence from which they were derived. This 25% had an average mass of 11.4 kDa, compared to the remaining 75% which had an average mass of 5.7 kDa. Several factors may contribute to this observation, some of which are independent of FAIMS. For example, these fragments themselves may be proteolytic cleavage products produced under normal homeostatic conditions as part of the cellular “degradome”,^62^ or despite the various precautions taken, introduced during the postmortem interval and sample handling. Central nervous system tissue is a rich source of signaling peptides known as “neuropeptides” that are commonly derived from much larger precursor proteins as well.^63^ By cross-referencing our proteoform identifications with an established neuropeptide database (NeuroPedia)^64^ we were able to identify several such neuropeptides including vasostatin-1, secretoneurin, cholecystokinin-58 desnonopeptide, neuropeptide Y, and many non-canonical sequence variants derived from known neuropeptide-producing genes. Outside of biological factors, certain instrument parameters can influence the observation of protein fragments as well. For example, low in-source “fragmentation” voltages (between 10-20 V) can typically be used to remove adducts and desolvate protein ions, while higher voltages can produce source-induced dissociation. However, the susceptibility of amide bonds to dissociation can vary widely, and as the size of a protein increases so does the likelihood of it containing labile amide bonds such as Xaa-Pro.^65–68^ As *b*- and *y*-ions may be produced inadvertently through this mechanism, even at low source voltages, we decided to eschew applying source voltage to reduce the chance of introducing fragment ions into the instrument. It is also possible proteins are susceptible to fragmentation events as they pass through the electric fields created in FAIMS. However, these fragments would not be expected to have the same mobility as the parent ion, with the caveat that this is likely dependent on the size of the fragment relative to the precursor ion as well.

Finally, it is worth pointing out a few factors that can bias towards the observation of smaller proteoforms (< ~15 kDa). First and foremost is the signal spreading that occurs as the size of a proteoform increases.^69^ This signal spreading can largely be attributed to the charge state envelope and isotopic distributions of each charge state. Specific to our experimental setup was the use of a size 3K MWCO filter as the final filtration step in our sample prep, which in this case was chosen to ensure smaller amyloid beta proteoforms could be efficiently captured if they were present. General instrumental factors that can create bias towards small proteoforms including the tuning of the quadruople, electrodynamic capture, and collisions with residual gas molecules in the Orbitrap cell that lead to quicker decay of the transient for larger molecules.^70^ Irrespective of these biases or the exact origin of the fragments, they can still provide insight into the proteome. These fragments are particularly useful in the context of neurodegenerative disease where disruptions in proteostasis are commonly linked to pathology.^71, 72^ Indeed, fragments of tau are often found to be neurotoxic and play a role in the progression of tauopathies such as Alzheimer’s disease.^23–26^ Generally, the fragments we observe are much larger than tryptic peptides and in several cases where there are many fragments for a particular protein, we observe considerable sequence coverage.

This is exemplified by examining the fragments providing coverage for the ~50 kDa tubulin alpha-1B chain (TUBA1B) and the ~70 kDa synapsin-1 (SYN1) in **Figure 9A** and **9B**, both large proteins that would otherwise be difficult to observe in their full-length form in TDP of a complex sample.

**Figure 9:**
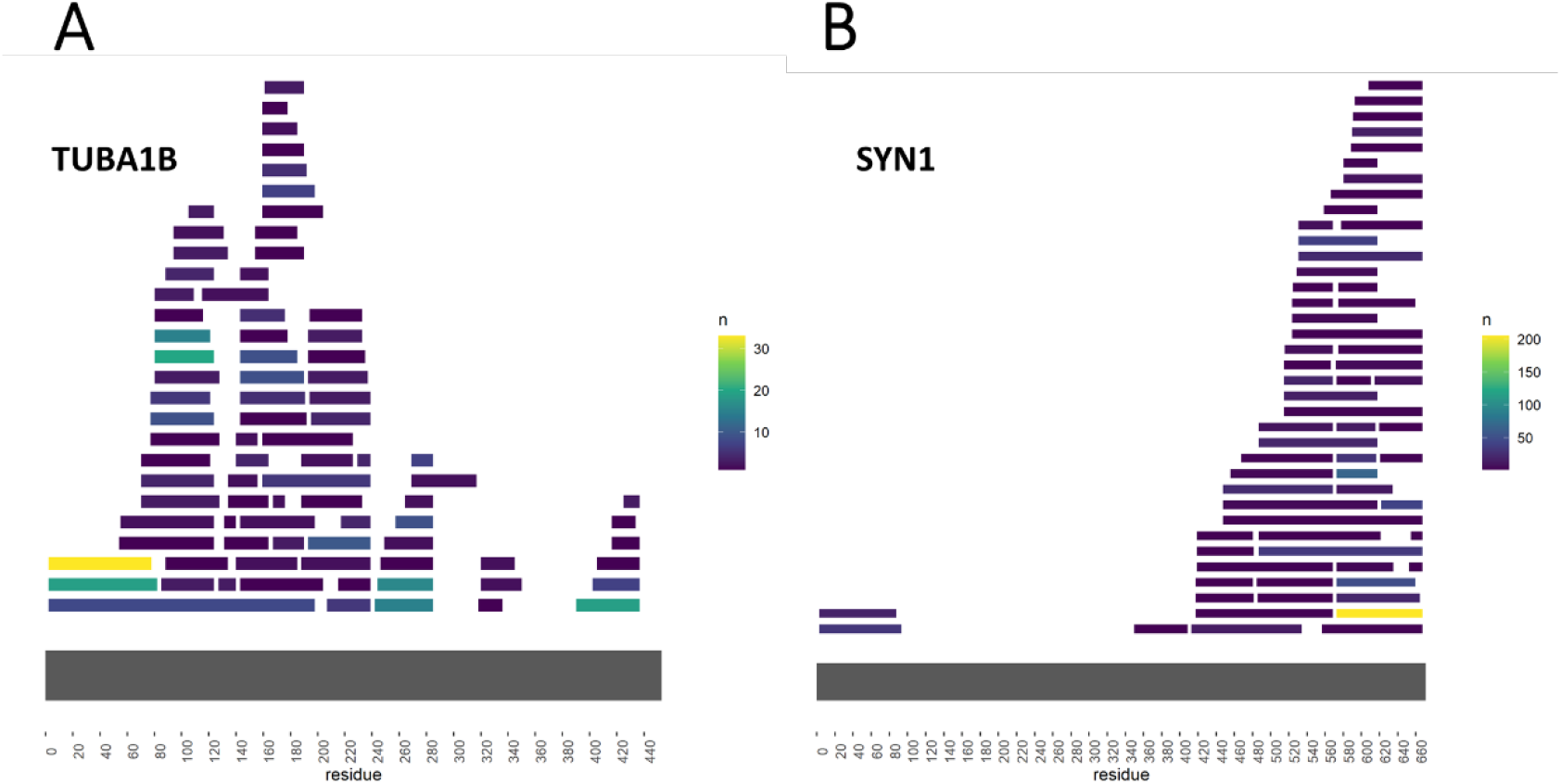
Sequence maps of (**A**) tubulin alpha-1B chain (TUBA1B) and (**B**) synapsin-1 (SYN1). Each colored segment represents a proteoform with a unique amino acid start and end site. Color scale represents the number of observed PrSMs that can be mapped to the corresponding segment. Unknown mass shifts were not utilized in this analysis.

### Identification of Proteoforms with Relevance to Neurodegenerative Disease

The utilization of FAIMS with our top-down analysis enabled identification of a number of unique Swiss-Prot splice variants and TrEMBL entries, with 267 unique splice variants and 96 TrEMBL entries (**Table S3**). Notably, we observe multiple PrSMs unambiguously identifying an alternative ORF isoform (A0A0D9SF30) for neural cell adhesion molecule 1 (NCAM1). We also observed proteoforms derived from genes that have known roles in several neurodegenerative diseases, in particular α-synuclein and PARK7. In both of these cases, the predominant proteoforms contain the full-length sequence. Interestingly, the majority of α-synuclein, β-synuclein, and (to a lesser extent) γ-synuclein PrSMs were found to contain a ~177 Da unknown mass shift near the C-terminus (full length α-synuclein spectrum with unknown modification shown in **Figure S4A**).This mass shift has been previously observed in open database searches and is generally found at Asp or Glu residues. ^73^ A potential explanation that closely matches the average and monoisotopic mass of the modification consists of one oxygen with three iron atoms as well as the loss of seven hydrogen atoms (based on Unimod accession #1971). Comparison of the isotopic peaks of a y^2+^_24_ fragment ion from α-synuclein (**Figure S4B**) with a simulated spectrum containing the aforementioned elements thought to belong to the unknown modification (**Figure S4C**) demonstrates the similarity between the two isotopic distributions, with many of the isotopic peaks aligning within 2 ppm of each other. It should also be pointed out that the left most isotopic peaks are unique to naturally occurring iron isotopes, and the absence of these peaks is readily apparent when the spectrum is simulated without containing the three iron atoms (**Figure S4D**), strongly suggesting this unknown modification is very likely composed of the proposed elements. Prior studies have also described the high affinity of α-synuclein for various metal ions, and the region we observe to contain this modification overlaps with residues known to be involved in binding - specifically the ^119^DPDNEA^124^ motif (**Figure S5**).^74–76^ Our data also suggests that the C-terminal regions of β-synuclein and γ-synuclein have similar roles in metal binding as multiple spectra with the same unknown mass shift were matched to similar sequences within these proteins. With regards to the mitochondrial protein PARK7, we noted a ~116.0 Da mass shift on its single active site Cys residue, C^106^ (**Figure S6**). One potential explanation with a similar delta mass is succinylation, a modification that is attributed to mitochondrial stress and forms due to Michael addition of fumarate onto a Cys thiol group.^77^

We also identified numerous fragments derived from non-canonical splice isoforms of tau protein known to be expressed in the human brain.^78^ Briefly, tau isoforms are defined by the number of N-terminal inserts due to alternative splicing of exon 2 and/or 3 (referred to as 0N, 1N, and 2N), and the number of microtubule-binding repeats due to alternative splicing of exon 10 (referred to as 3R or 4R).^79^ It’s also worth noting that endogenous fragments of tau are a common finding in human brain tissue and CSF fluid, and differential fragmentation of tau may play a role in Alzheimer’s disease progression.^80, 81^ We observed several fragments that could unambiguously distinguish 0N and 1N tau (**Figure 10A**, **S7**, and **S8**). The fact that we observe high spectral counts for 0N (86 PrSMs) and 1N (91 PrSMs), but do not observe any PrSMs for 2N, is aligned with previous quantitative immunoblotting that found 0N and 1N to be the dominant forms and make up ~91% of total tau while 2N comprised only ~9%.^82^ With regards to unambiguous assignment of the microtubule-binding repeats we observed spectra that could be assigned to both 3R (**Figure 10A** and **S9**)and 4R tau (**Figure 10A** and **S10**) as well.

**Figure 10:**
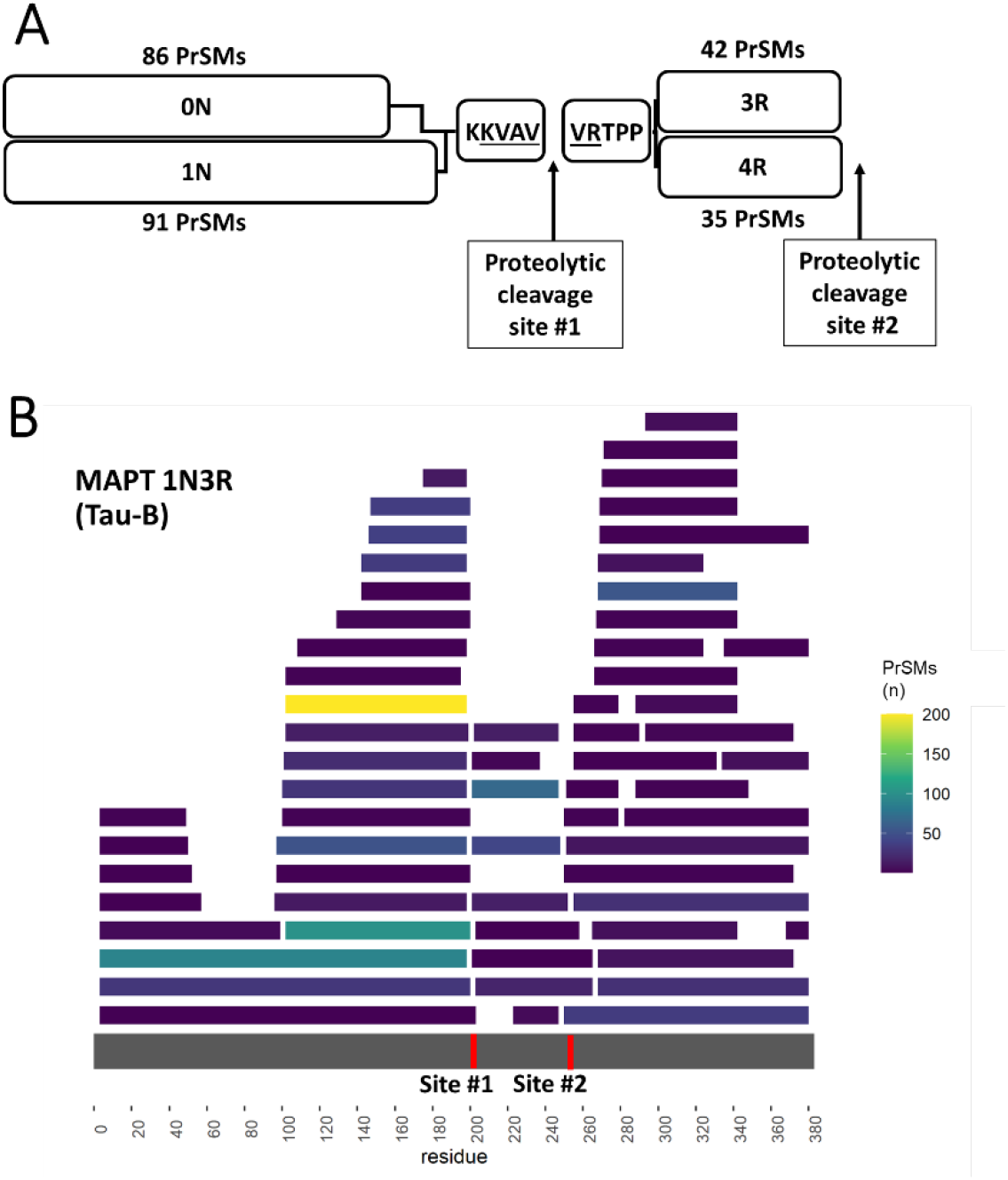
(**A**) Schematic representation of selected tau fragments with the number of PrSMs that unambiguously identify 0/1N or 3/4R tau. 0N and 1N fragments are being shown as sharing the same KKVAV residues at their C-terminus, while 3R and 4R proteoforms share VRTPP at their N-terminus. Residues underlined are the KVAVVR hexapeptide sequence in the second-proline rich region of tau which is cleaved at proteolytic cleavage site #1. The 3R and 4R hexapeptide motifs are not shown but are both cleaved at proteolytic cleavage site #2. (**B**) Sequence coverage map of MAPT 1N3R (Tau-B) showing all proteoforms that could be mapped to this isoform with a unique amino acid start and end site. Red lines at the bottom represent the positions of proteolytic cleavage sites #1 and #2. Color scale represents the number of observed PrSMs that can be mapped to the corresponding segment. Unknown mass shifts were not utilized in this analysis.

Parsimonious inference allowed us to conclude that brain tau is represented mostly by mixture of 1N3R (Tau-B) followed by 0N4R (Tau-D) splice variants, which is in agreement with previous observations.^82^’ ^83^ While other splice isoforms also had unique fragments, the low number of spectral matches (<4) did not allow us to confidently conclude about their existence.

Surprisingly, N-termini of the predominant R-domain containing fragments are a continuation from the C-termini of the predominant N-domain containing fragments, or in other words they appear to be linked fragments produced from a possible proteolytic cleavage event (referred to as site #1 in **Figure 10A**). Furthermore, the C-termini of the 3R and 4R domain-containing proteoforms share the same cleavage site (referred to as site #2 in **Figure 10A**) even though the sequences themselves are different due to splice variation of exon 10. Even more surprising, all of these cleavage sites lie within three well described hexapeptide motifs.^84–90^ **Figure 10B** demonstrates all of the unique fragments we identified from our datasets that could be mapped to the 1N3R tau isoform, as well as the location of the cleavage sites. Cleavage site #1 lies within the second proline-rich region of tau and divides the KVAVVR hexapeptide sequence which is known to provide one of the strongest binding sites for microtubules (**Figure 10B**).^84–86^ Cleavage site #2 lies within the 3R and 4R hexapeptide motifs VQIVYK and VQIINK, which have been demonstrated to drive aggregation of tau (**Figure 10B**).^87–90^ Interestingly, many of the fragments shown in **Figure 10B** begin or end within proximity of these two cleavage sites. Each of these hexapeptide motifs are known to form β-structure, often in the form of β-hairpins which underlie the aggregation involved in various neurodegenerative diseases.^84, 88, 91^ Intriguingly, these observations highlight the possibility of a shared proteolytic degradation pathway among tau isoforms that is potentially capable of disrupting their aggregation-seeding regions.^25, 80, 81^

Lastly, we found numerous spectral matches for Aβ that could only be observed exclusively by using FAIMS. These Aβ proteoforms, which typically require special fractionation or handling techniques in order to improve recovery due to their hydrophobic and aggregation prone-properties,^9, 16, 92, 93^ were observed intact using FAIMS within the −40 to −50 CV range. This includes the canonical Aβ_1-42_ and Aβ_1-40_ (**Figure 11A** and **11B**, respectively), as well as several N-terminally truncated forms (specifically Aβ_2-42_ and Aβ_4-42_, **Figure 11C** and **11D**).

**Figure 11:**
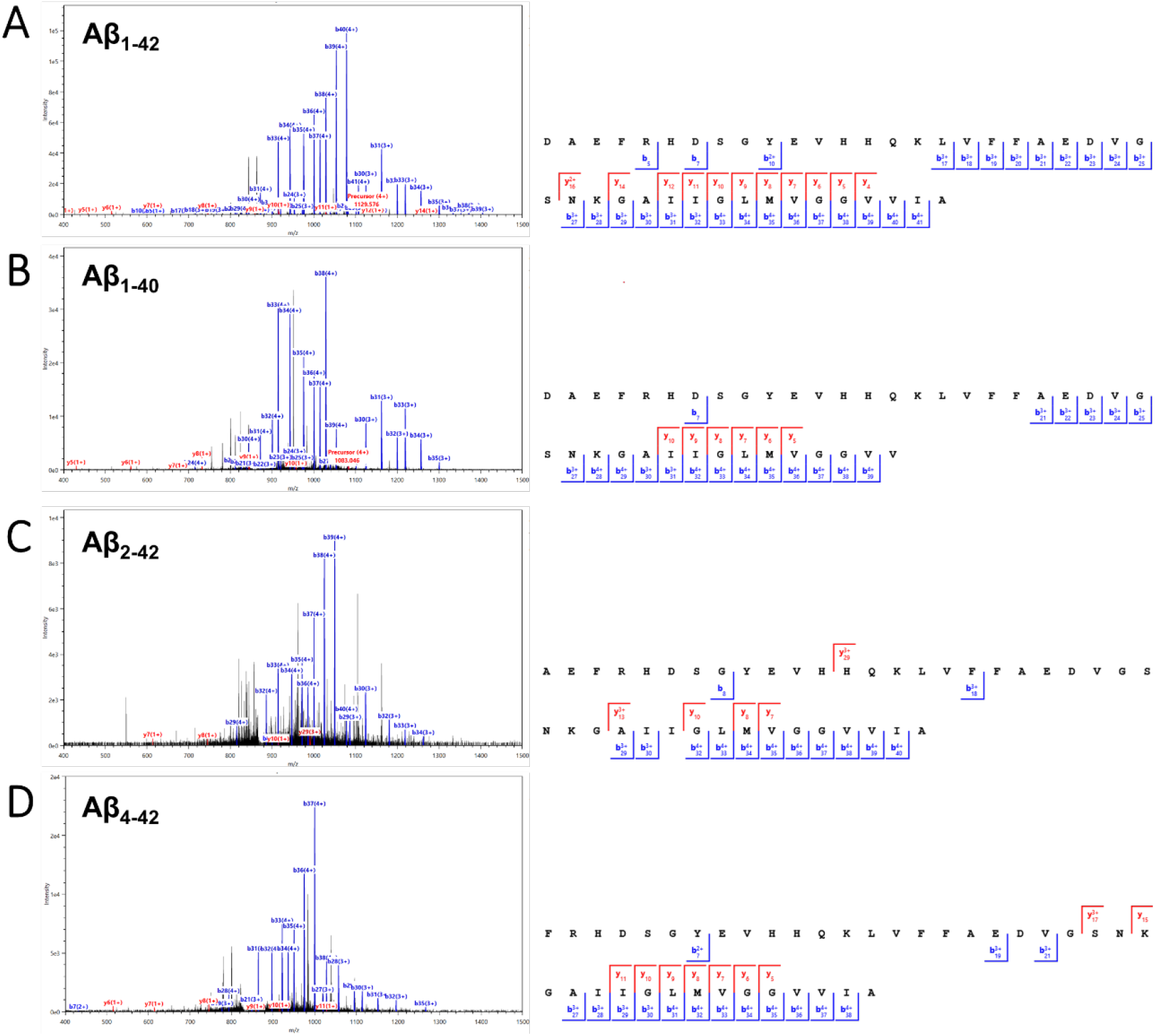
Representative MS2 spectra and MS2 fragment ion coverage maps with matched *b* and *y* ions of the 5+ charge state of four Aβ proteoforms: (**A**) Aβ_1-42_ from 903.663 *m/z* precursor with 68.3% sequence coverage (43 PrSMs). (**B**) Aβ_1-40_ from 866.4374 m/z precursor with 48.7% sequence coverage (6 PrSMs). (**C**) Aβ_2-42_ from 880.6570 m/z precursor with 40.0% sequence coverage (3 PrSMs). (**D**) Aβ_4-42_ from 840.6410 *m/z* precursor with 44.7% sequence coverage (4 PrSMs).

It is worth noting that Aβ proteoforms with various N- and C-terminal cleavages are notoriously difficult to identify in bottom-up analyses due to the extreme hydrophobicity of the tryptic products, which highlights an advantage of intact protein analysis. We believe the FAIMS-TDP methodology offers a robust way for achieving intact identification of these Aβ proteoforms, allowing for facile determination of the different combinations of N- and C-terminal cleavage products that exist.

## Conclusions

We have described how nano-LC reverse-phase separation of a highly complex sample containing proteins over a wide mass range, can benefit from the implementation of gas-phase fractionation through FAIMS in the context of top-down mass spectrometry. FAIMS was demonstrated to impact the transmission of proteoforms by size and/or charge, reducing MS^1^ complexity and allowing greater depth of coverage of the proteome. FAIMS at a single CV (−50) enabled identification of 1,833± 17 unique proteoforms on average, more than double compared to without FAIMS (754 ± 35). Addition of FAIMS did not result in deterioration of quantification reproducibility and remained near 20% RSD, which is comparable with label-free bottom-up approaches. Decreasing CV of FAIMS was noted to increase the molecular mass of ion species being transmitted to the instrument (median MW of ~5 kDa at −50 CV vs ~15 kDa at −20 CV), and modulation of CV was also observed to differentially influence the transmission a proteoform’s charge state envelope. We also defined optimal combinations of CVs that could produce the largest number of proteoforms/genes (−50, −40, −35 CV) or proteome sequence coverage (−50, −40, and −30 CV). It is worth pointing out that our work only explored a relatively small CV range which we felt best benefited TDP of a complex sample. Based on our results here, future mass analyzer instrumentation compatible with FAIMS that improve on the acquisition of larger proteoforms may benefit from exploring CVs below −20, extending into the positive voltage range.

Our TDP workflow allowed us to identify and characterize unique proteoforms derived from genes with known roles in neurodegenerative diseases. This included determination of the composition of an unknown mass shift present near the iron-binding domain of α-synuclein proteins, which was used to pinpoint the locations of potential iron-binding domains in β- and γ-synuclein as well. PARK7 was also found to be modified with a succinyl group on its active site Cys residue. We were able to unambiguously distinguish tau fragments corresponding to 0N, 1N, 3R, and 4R splice variant isoforms individually, and describe new proteolytic cleavage sites located within or near several aggregation-seeding hexapeptide repeats. Finally, FAIMS enabled identification of several intact Aβ proteoforms, including the aggregation-prone Aβ_1-42_, without the need for complex fractionation or purification techniques. In summary, we believe that addition of FAIMS to discovery top-down proteomics will provide a robust and reproducible method for increasing the proteome available for detection and characterization.

## Acknowledgments

The authors would like to thank Tao Liu and Richard Smith for advice and helpful discussions. This work was supported by U01 AG061356 (P.L.D.J.) and R01 AG015819 (D.A.B.). A portion of the research was performed using EMSL (grid.436923.9), a DOE Office of Science User Facility sponsored by the Biological and Environmental Research program.

