## Supplemental Information for "Enhancing top-down proteomics of brain tissue with FAIMS"

*To whom correspondence should be addressed

**Table of Contents:**


| Title | Page |
|---|---|
| Table S1–Table of proteoform metrics | 3 |
| Figure S1–Reproducibility of FAIMS and No FAIMS TDP Data | 4-5 |
| Figure S2–Overlap coefficient heatmap of genes | 5 |
| Figure S3–Consecutive $MS^1$ scans with median charge states of ubiquitin | 6 |
| Table S2–Table of proteoform characteristics | 7 |
| Table S3–Table of unique Swiss-Prot splice variants and TrEMBL entries | 7 |
| Figure S4–Characterization of unknown modification on α-synuclein | 8 |
| Figure S5–$MS^2$ fragment coverage map of α-synuclein | 9 |
| Figure S6–$MS^2$ spectrum and coverage map of succinylated PARK7 | 10 |
| Figure S7–$MS^2$ spectrum and coverage map of Tau(0N-R) fragment | 11 |
| Figure S8–$MS^2$ spectrum and coverage map of Tau(1N-R) fragment | 12 |
| Figure S9–$MS^2$ spectrum and coverage map of Tau(-N3R) fragment | 13 |
| Figure S10–$MS^2$ spectrum and coverage map of Tau(-N4R) fragment | 13 |

| Condition (n) | Mean Proteoforms Observed (+/-SD) | Unique Proteoforms | Unique Genes | Proteome Sequence Coverage (#AA) |
|---|---|---|---|---|
| -50 CV (3) | 1833 ± 17 | 2564 | 530 | 43,437 |
| -45 CV (3) | 1709 ± 84 | 2449 | 522 | 47,621 |
| -40 CV (3) | 1442 ± 80 | 2094 | 496 | 49,720 |
| -35 CV (3) | 1147 ± 25 | 1656 | 431 | 48,668 |
| -30 CV (3) | 781 ± 8 | 1092 | 341 | 41,093 |
| -25 CV (3) | 403 ± 15 | 563 | 206 | 28,398 |
| -20 CV (3) | 200 ± 2 | 288 | 117 | 18,243 |
| All FAIMS (21) | - | 5165 | 799 | 88,046 |
| No FAIMS (3) | 754 ± 35 | 1073 | 293 | 29,359 |

**Table S1:** Table of values used in Figure 2; including mean proteoforms observed, total unique proteoforms, total unique genes, and proteome sequence coverage (measured with number of amino acids, #AA) found across the specified number of replicates and conditions.

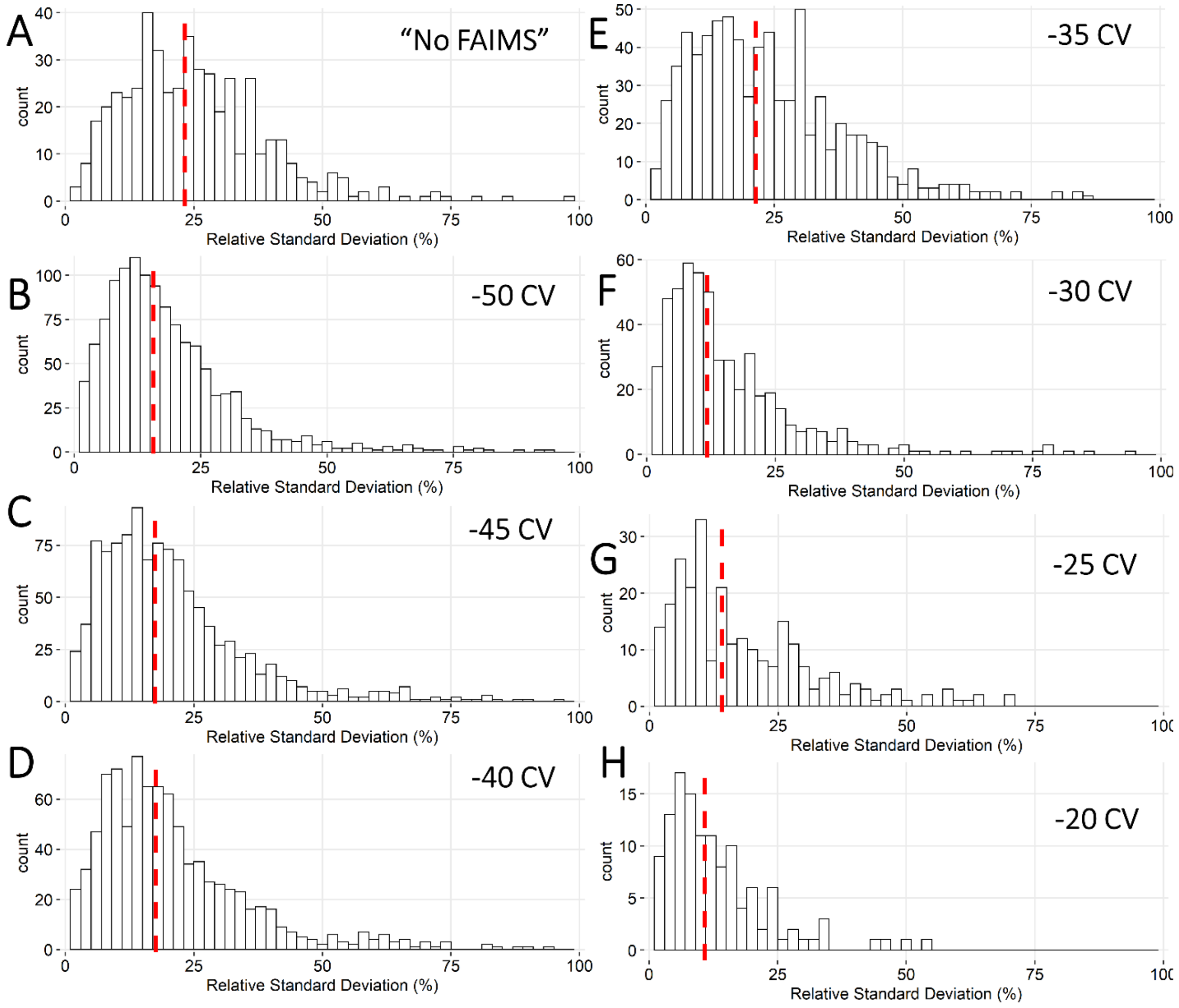


**Figure S1**. Reproducibility analysis of replicates for all CVs of FAIMS as well as "No FAIMS" datasets. For each condition, proteoforms that were identified in all three replicates were binned in steps of 2 by the relative standard deviation (RSD) of their feature intensities across the replicates. Red dotted lines indicate the median of the distribution. Feature intensities within each replicate were normalized via mean normalization. Outliers beyond 100% RSD are not displayed on these histograms. (**A**) Histogram of the RSDs for the feature intensity of all proteoforms identified across the three replicates with "No FAIMS". 492 proteoforms were able to be quantified across all three replicates, producing a median feature intensity RSD of 23.2%. (**B**) Histogram of the RSDs for the feature intensity of all proteoforms identified across the three replicates at -50 CV with FAIMS. 1,228 proteoforms were able to be quantified across all three replicates, producing a median feature intensity RSD of 15.6%. (**C**) Histogram of the RSDs for the feature intensity of all proteoforms identified across the three replicates at -45 CV with FAIMS. 1,106 proteoforms were able to be quantified across all three replicates, producing a median feature intensity RSD of 17.5%. (**D**) Histogram of the RSDs for the feature intensity of all proteoforms identified across the three replicates at -40 CV with FAIMS. 909 proteoforms were able to be quantified across all three replicates, producing a median feature intensity RSD of 17.6%. (**E**) Histogram of the RSDs for the feature intensity of all proteoforms identified across the three replicates at -35 CV with FAIMS. 738 proteoforms were able to be quantified across all three replicates, producing a median feature intensity RSD of 21.4%. (**F**) Histogram of the RSDs for the feature intensity of all proteoforms identified across the three replicates at -30 CV with FAIMS. 530 proteoforms were able to be quantified across all three replicates, producing a median feature intensity RSD of 11.7%. (**G**) Histogram of the RSDs for the feature intensity of all proteoforms identified across the three replicates at -25 CV with FAIMS. 268 proteoforms were able to be quantified across all three replicates, producing a median feature intensity RSD of 14.1%. (**H**) Histogram of the RSDs for the feature intensity of all proteoforms identified across the three replicates at -20 CV with FAIMS. 126 proteoforms were able to be quantified across all three replicates, producing a median feature intensity RSD of 10.8%.

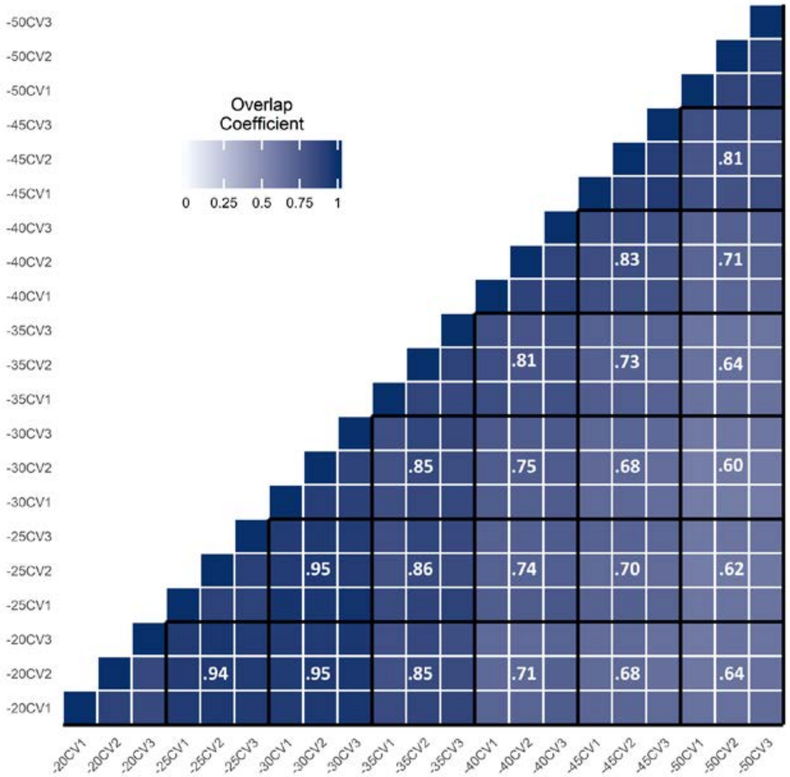


**Figure S2**: Heatmap generated in R comparing the overlap coefficients of each CV replicate based on genes identified with TopPIC. White lines separate replicates while black lines separate different CVs. Mean overlap coefficients of each CV's replicates are shown in the middle block of the CVs being compared. White text color is used on overlap coefficients >= 0.5 and black text color < 0.5 to improve visibility.

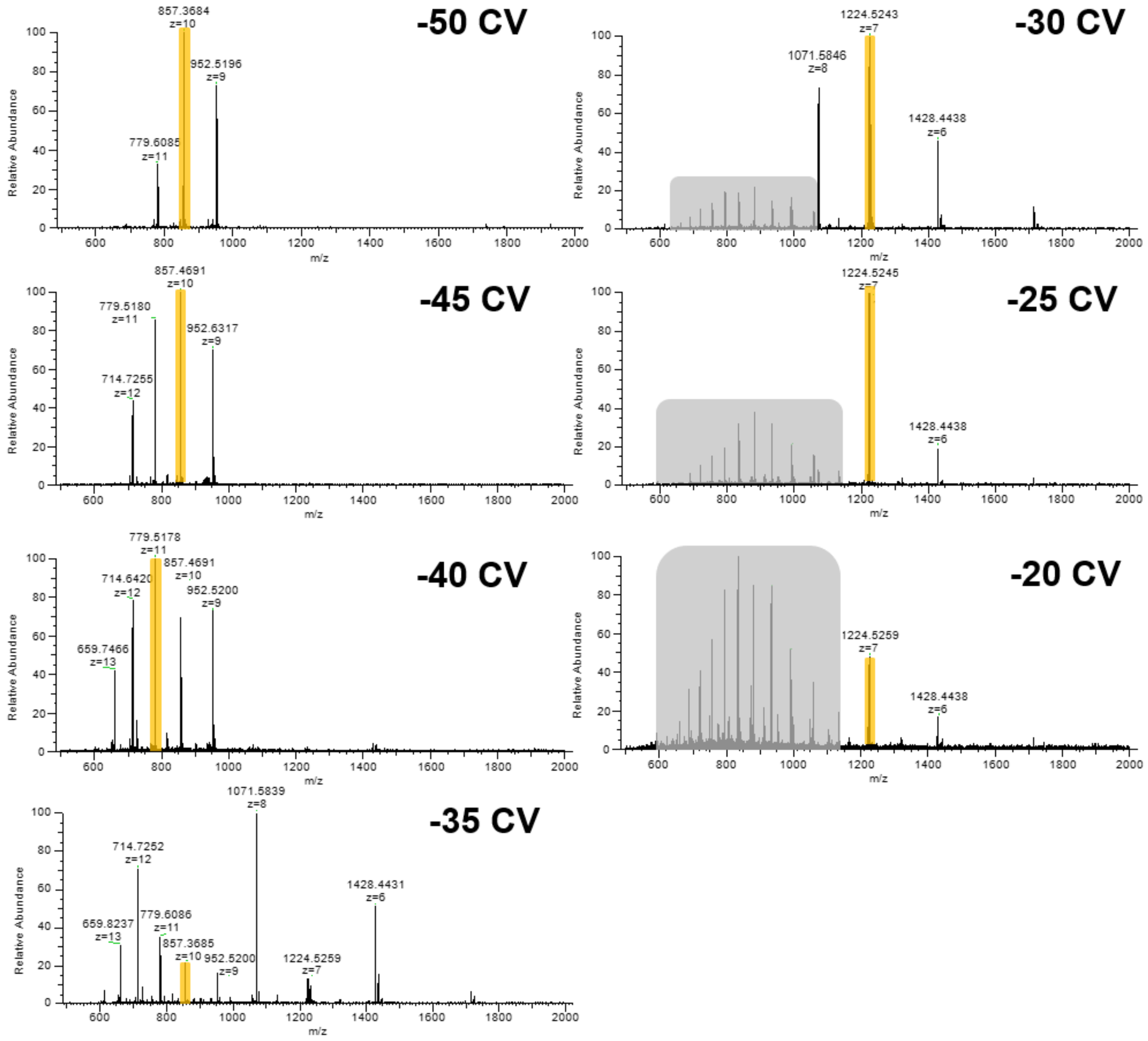


**Figure S3:** Consecutive MS$^1$ scans of ubiquitin across the -50 to -20 CV range in steps of 5 V, demonstrating how the median charge state tracks the ubiquitin charge state envelope with respect to CV. Orange highlighting indicates location of the median charge state rounded to the nearest whole number. Peaks with charge and mass labels correspond to ubiquitin, while gray highlighting represents a separate protein species present in the spectrum.

| Relationship | Average Precursor Mass | Average % Basic Residues [K,R,H] | Mean % Acidic Residues [D,E] | Aliphatic Index | n |
|---|---|---|---|---|---|
| Inverse | 11,440 ± 2800 Da | 18.4 ± 4.4% | 12.1 ± 5.4% | 70.4 ± 21.0 | 39 |
| Direct | 7770 ± 2360 Da | 18.6 ± 5.9% | 13.8 ± 6.8% | 71.7 ± 23.8 | 177 |
| Neutral | 9240 ± 4430 Da | 18.0 ± 5.0% | 11.9 ± 7.5% | 71.0 ± 21.3 | 40 |

**Table S2**: Table of different primary sequence properties for proteoforms found within a 20-30 voltage range using FAIMS sorted by the differential transmission of their charge states as CV is modulated (Inverse, Direct, or Neutral).

| Condition (n) | Unique Swiss-Prot Splice Variants | Unique TrEMBL Entries |
|---|---|---|
| -50 CV (3) | 81 | 22 |
| -45 CV (3) | 79 | 34 |
| -40 CV (3) | 83 | 43 |
| -35 CV (3) | 111 | 36 |
| -30 CV (3) | 89 | 19 |
| -25 CV (3) | 57 | 2 |
| -20 CV (3) | 27 | 0 |
| All FAIMS (21) | 267 | 96 |
| No FAIMS (3) | 69 | 15 |

**Table S3**: Table listing the number of unique Swiss-Prot splice variants and TrEMBL entries found within different FAIMS and “No FAIMS” datasets.

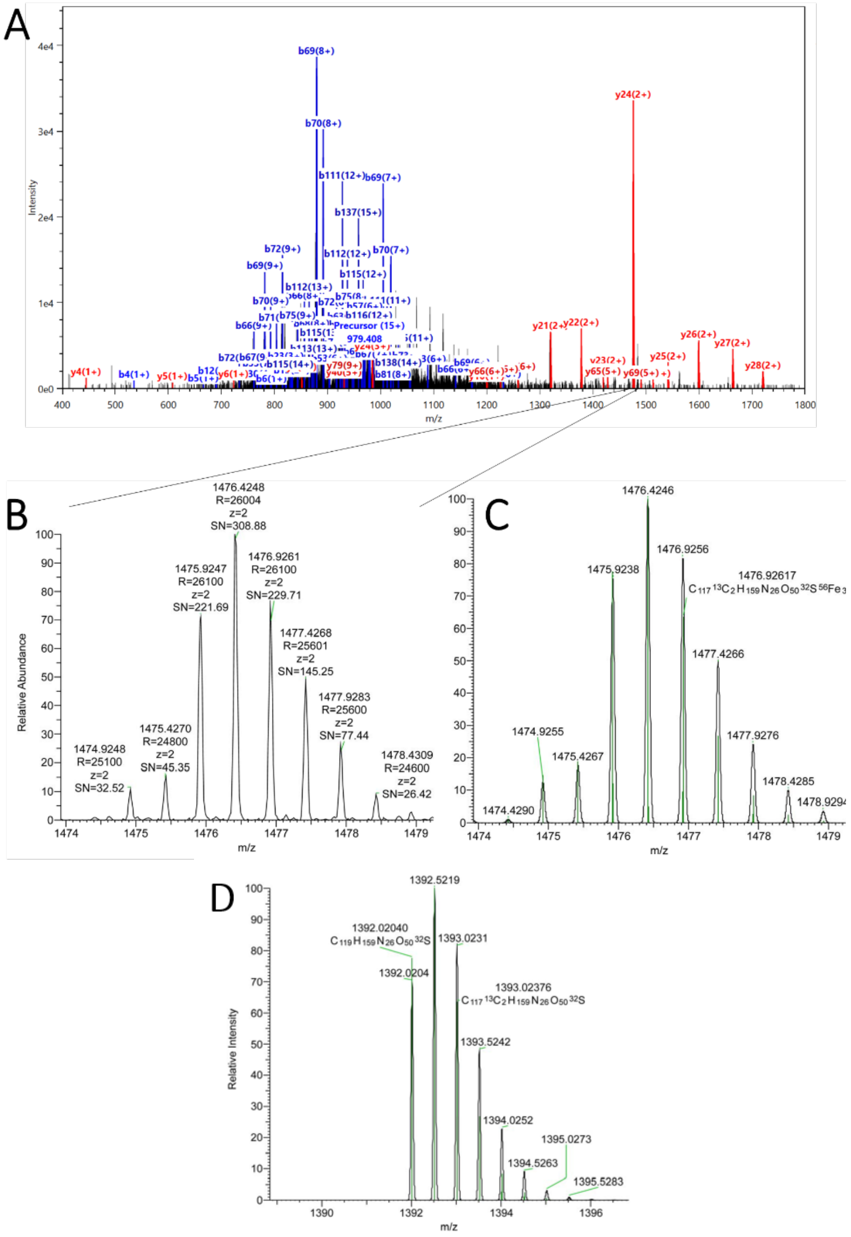


**Figure S4**: (**A**) Representative MS$^2$ spectrum with matched *b* and *y* ions of the 15+ charge state precursor (979.408 m/z) of full-length a-synuclein with N-terminal acetylation and an unknown mass shift of 176.744957 on D121. (**B**) Close-up view of the $y_{24}^{2+}$ containing the unknown mass shift from. (**C**) Simulated isotopic distribution of the $y_{24}^{2+}$ ion with the addition of three iron atoms, one oxygen atom, and removal of seven hydrogen atoms ((PVDPD[176.744957]NEAYEMPSEEGYQDYEPEA). Empirical formula used to generate simulated spectrum for the $y_{24}^{2+}$ peptide with unknown modification was $C_{119}H_{157}N_{26}O_{50}SFe_3$. (**D**) Simulated isotopic distribution of the $y_{24}^{2+}$ ion without the three iron atoms demonstrating the diagnostic loss of the iron isotopes. Empirical formula used to generate this simulated spectrum was $C_{119}H_{157}N_{26}O_{50}S$. Simulated spectra were generated using Thermo Fisher’s FreeStyle 1.7 SP1.

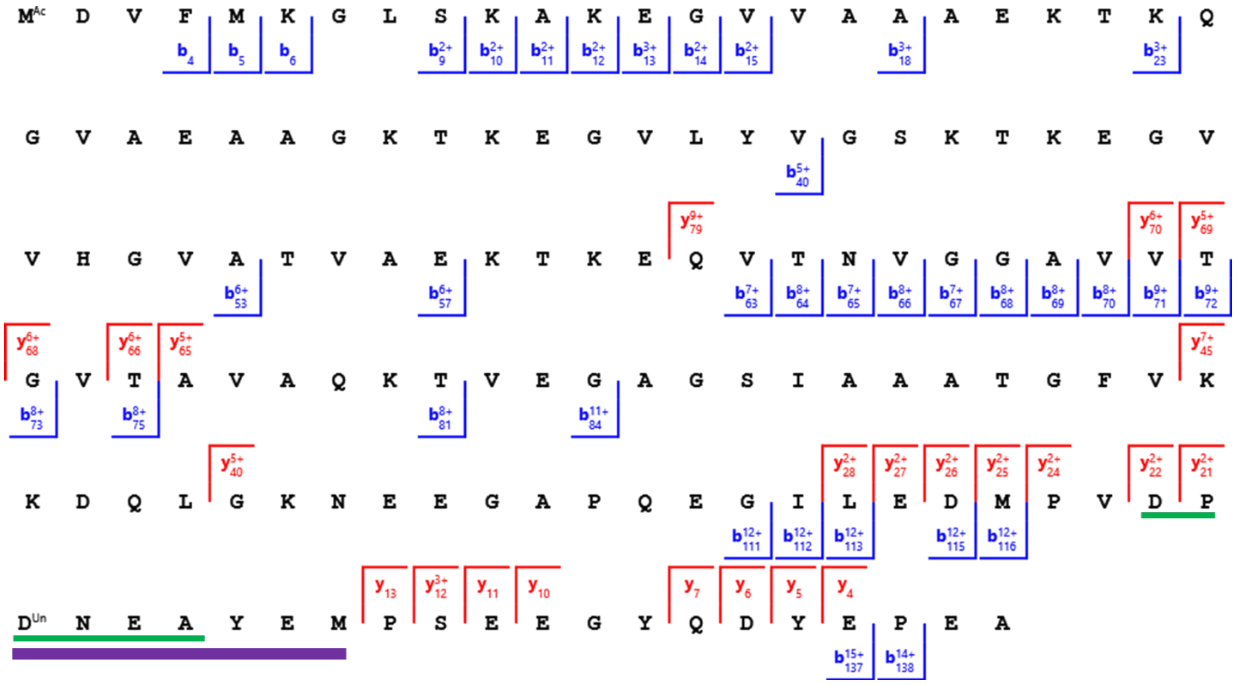

**Figure S5:** MS$^2$ fragment ion coverage map (36.0% coverage) of full-length alpha-synuclein derived from the 15+ charge state (979.408 m/z) MS$^2$ spectrum with matched *b* and *y* ions, including N-terminal acetylation and addition of a 176.744957 Da unknown modification onto amino acid D$^{121}$. Residues underlined in green ($^{119}$DPDNEA$^{124}$) represent previously studied iron-binding motifs from literature. Residues underlined in purple are the potential amino acids that the modification may be located at based on the *b* and *y*-ions from the spectrum in **Figure 10A**.

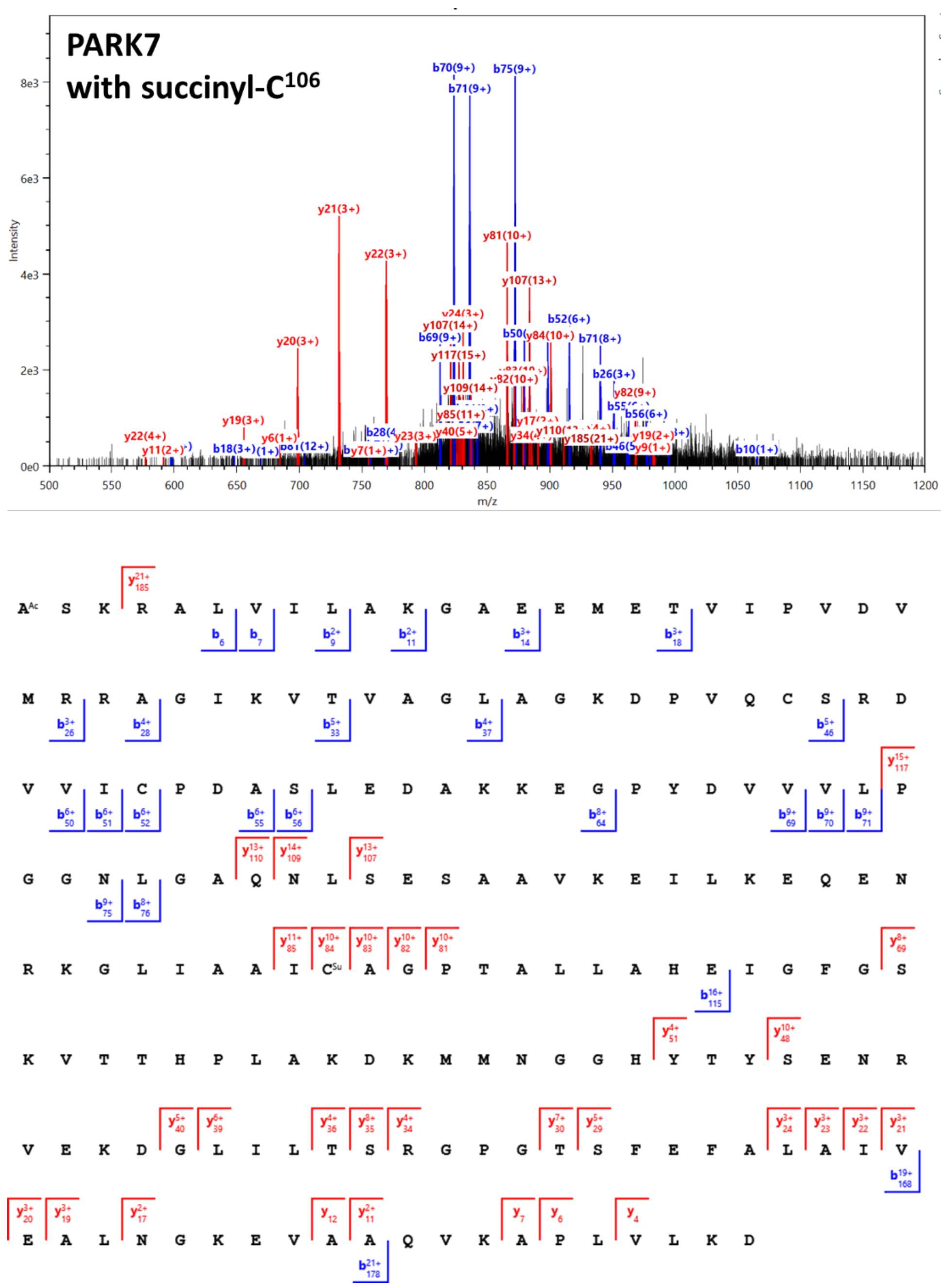


**Figure S6**: Representative $MS^2$ spectrum with matched *b* and *y* ions of the 25+ charge state precursor (797.2243 m/z) of full-length PARK7 with N-terminal acetylation and a succinyl (116.010959, C[4]O[4]H[4]) modification on $C^{106}$, and $MS^2$ fragment ion coverage map (29.8% coverage).

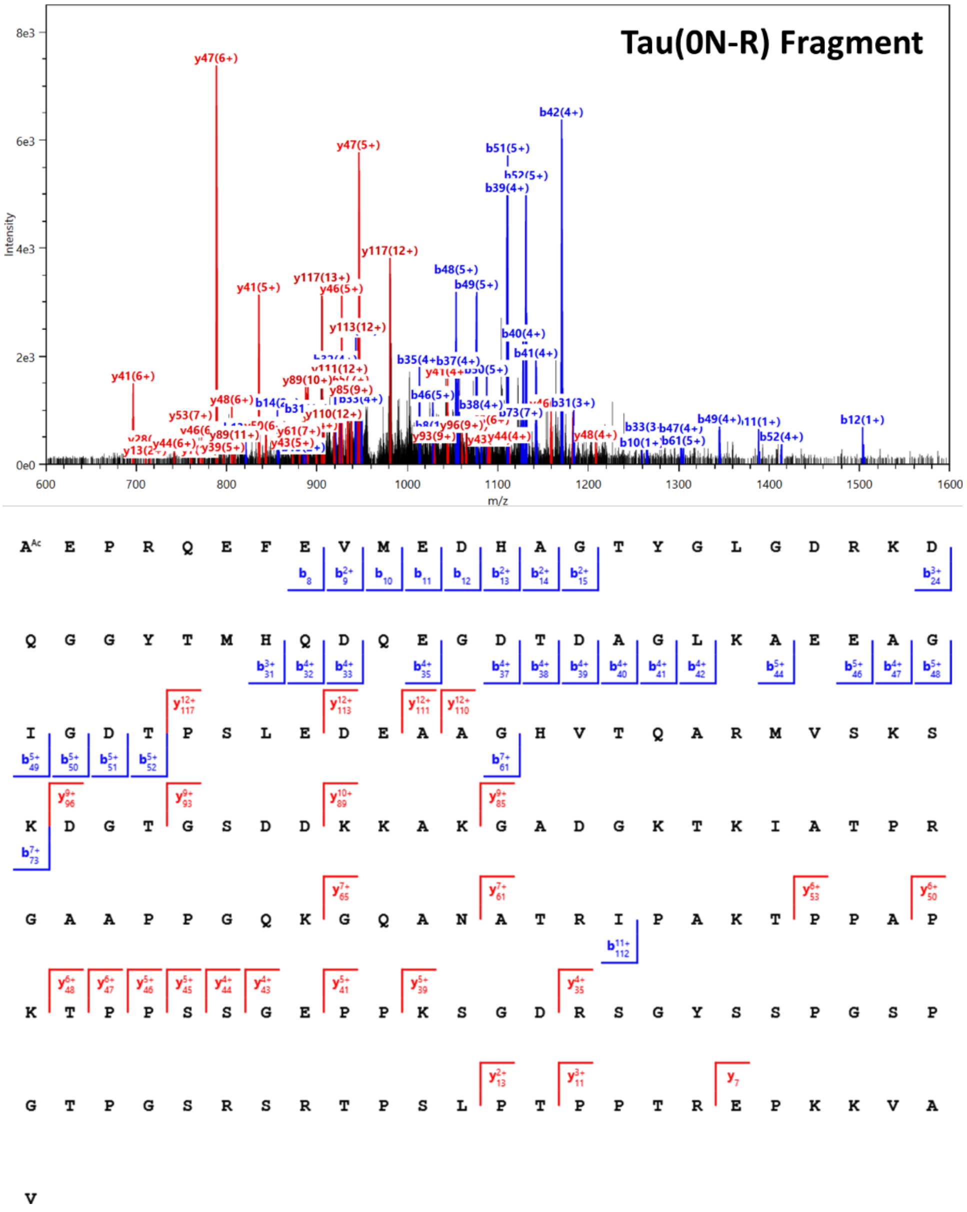


**Figure S7**: Representative MS$^2$ spectrum with matched *b* and *y* ions of the 18+ charge state precursor (967.7594 m/z) of a fragment of Tau(0N-R) with N-terminal acetylation, as well as MS$^2$ fragment ion coverage map (31.0% coverage) derived from the MS$^2$ spectrum.

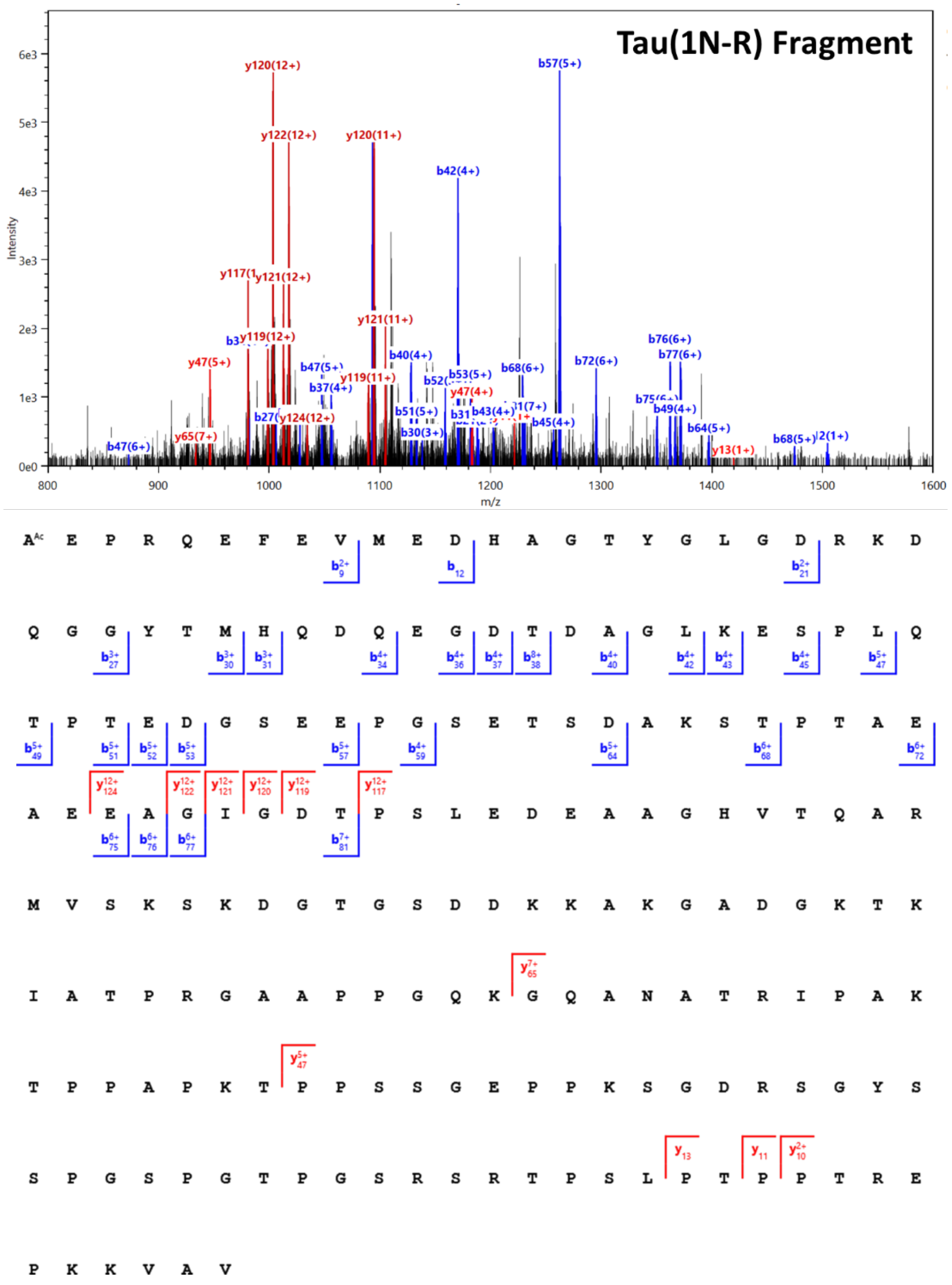


**Figure S8**: Representative MS$^2$ spectrum with matched *b* and *y* ions of the 19+ charge state precursor (1072.5787 m/z) of a fragment of Tau(1N-R) with N-terminal acetylation, as well as MS$^2$ fragment ion coverage map (18.3 % coverage) derived from the MS$^2$ spectrum.

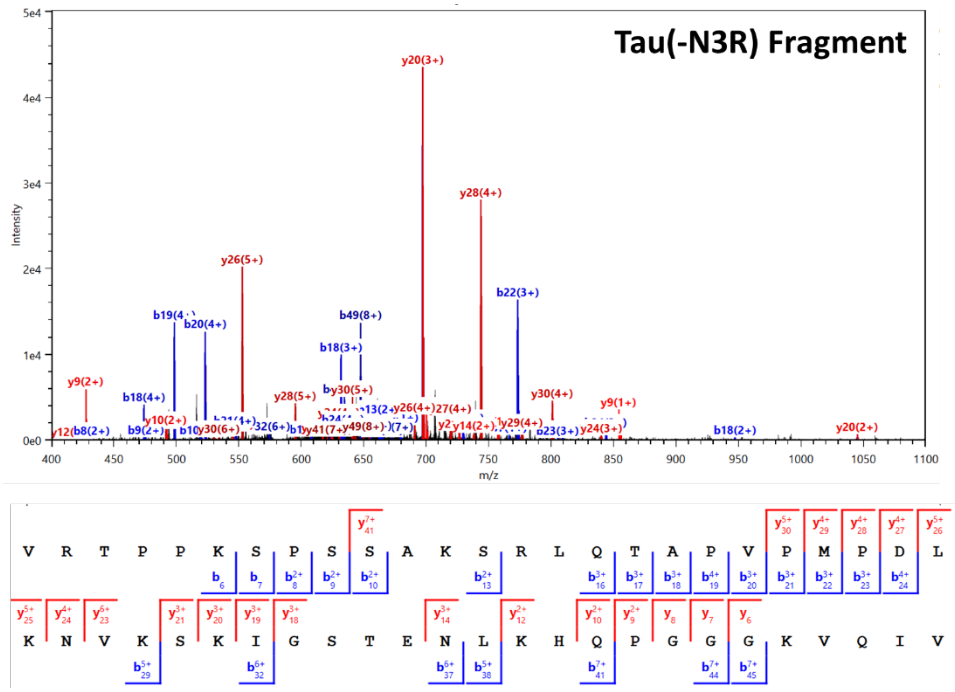


**Figure S9**: Representative MS$^2$ spectrum with matched *b* and *y* ions of the 9+ charge state precursor (588.4432 m/z) of a fragment of Tau(-N3R), as well as MS$^2$ fragment ion coverage map (67.3 % coverage) derived from the MS$^2$ spectrum.

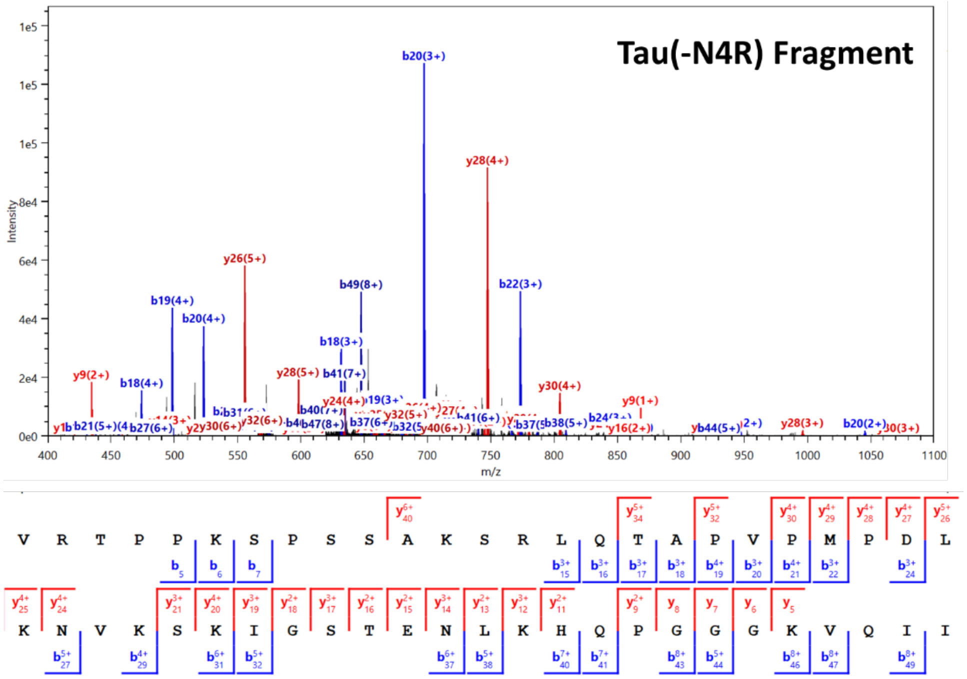


**Figure S10**: Representative MS$^2$ spectrum with matched *b* and *y* ions of the 9+ charge state precursor (590.0018 m/z) of a fragment of Tau(-N4R), as well as MS$^2$ fragment ion coverage map (75.5 % coverage) derived from the MS$^2$ spectrum.
